# PplD is a de-N-acetylase of the cell wall linkage unit of streptococcal rhamnopolysaccharides

**DOI:** 10.1101/2021.09.23.461590

**Authors:** Jeffrey S. Rush, Prakash Parajuli, Alessandro Ruda, Jian Li, Amol A. Pohane, Svetlana Zamakhaeva, Mohammad M. Rahman, Jennifer C. Chang, Artemis Gogos, Cameron W. Kenner, Gérard Lambeau, Michael J. Federle, Konstantin V. Korotkov, Göran Widmalm, Natalia Korotkova

## Abstract

The cell wall of the human bacterial pathogen Group A Streptococcus (GAS) consists of peptidoglycan decorated with the Lancefield group A carbohydrate (GAC). GAC is a promising target for the development of GAS vaccines. In this study, employing chemical, compositional, and NMR methods, we show that GAC is attached to peptidoglycan via glucosamine 1-phosphate. This structural feature makes the GAC-peptidoglycan linkage highly sensitive to cleavage by nitrous acid and resistant to mild acid conditions. Using this characteristic of the GAS cell wall, we identify PplD as a protein required for deacetylation of linkage *N*-acetylglucosamine (GlcNAc). X-ray structural analysis indicates that PplD performs catalysis via a modified acid/base mechanism. Genetic surveys *in silico* together with functional analysis indicate that PplD homologs deacetylate the polysaccharide linkage in many streptococcal species. We further demonstrate that introduction of positive charges to the cell wall by GlcNAc deacetylation protects GAS against host cationic antimicrobial proteins.

## Introduction

In Gram-positive bacteria, a cell wall outside the plasma membrane determines shape, ensures integrity, and protects the cell from environmental stresses and host immune defense mechanisms. In many streptococci, lactococci, and enterococci, the cell wall consists of multiple layers of peptidoglycan covalently modified with rhamnose (Rha)-containing polysaccharides ^1^. These essential polysaccharides are functional homologs of the canonical poly(glycerol-phosphate) and poly(ribitol-phosphate) wall teichoic acids (WTAs) expressed by *Bacillus subtilis* and *Staphylococcus aureus* ^2^, and non-canonical choline-containing WTAs produced by the *Streptococcus mitis* group ^3^.

*Streptococcus pyogenes* (Group A Streptococcus, GAS) is a human pathogen of global significance for which a vaccine is not currently available ^4^. The key antigenic surface polymer of GAS, the Lancefield group A carbohydrate (GAC), is composed of a repeating →3)α-Rha(1→2)α-Rha(1→ disaccharide backbone with *N*-acetyl-β-D-glucosamine (GlcNAc) side-chain modifications attached to the 3-position of the α-1,2-linked Rha ^5^. GAC has attracted significant interest from the scientific and medical communities as a vaccine candidate due to its uniform and invariant expression in all GAS serotypes and the absence of the constitutive component of GAC, Rha, in humans ^6-11^. Furthermore, in proof-of-principle mouse immunization studies, protein-conjugated GAC vaccines demonstrated efficacy against GAS strains ^6,7^. To provide well-defined GAC for immunological studies and vaccine design, knowledge of the structure and biogenesis of GAC is essential ^9^. The molecular mechanism of GAC biosynthesis was proposed based on genetic and biochemical studies of two genomic loci encoded by *gacABCDEFGHIJKL* and *gacO* ^8,12-16^. GAC biosynthesis is initiated on the inside of the plasma membrane on a lipid carrier, undecaprenyl phosphate (Und-P), by the action of the UDP-GlcNAc:Und-P GlcNAc-1-phosphate transferase, GacO. After assembly of the polyrhamnose backbone by the sequential activity of rhamnosyltransferases GacB, GacC, GacG and GacF, the nascent polysaccharide is then translocated across the cell membrane via the ABC transporter GacDE (Supplementary Fig. 1). Enzymatic machinery composed of GacI, GacJ, GacK and GacL catalyzes the addition of the GlcNAc side-chains which are then partially decorated with glycerol phosphate (GroP) moieties by GacH ^15^. Finally, the LytR-CpsA-Psr (LCP) phosphotransferases transfer GAC from the lipid carrier to peptidoglycan (Fig. 1 **a**).

**Fig. 1.**
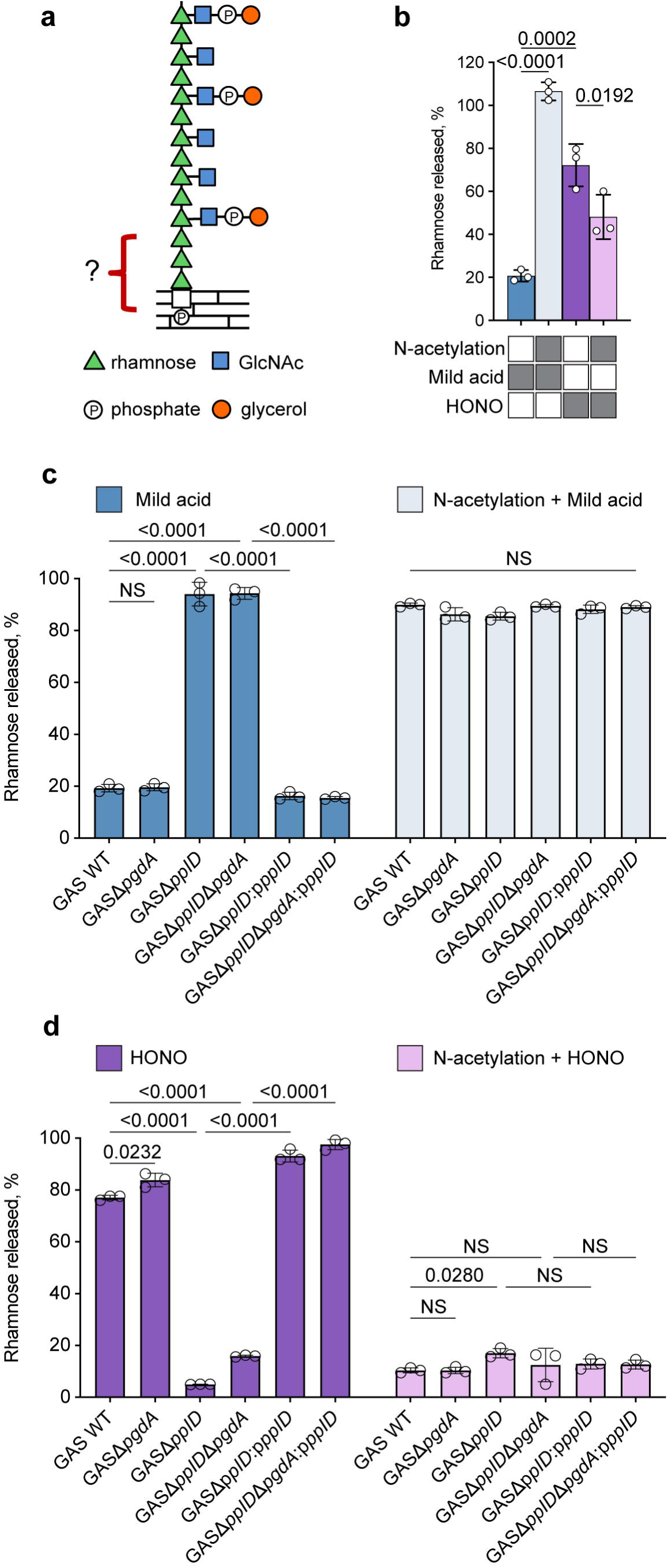
Analysis of GAC released from the GAS cell wall by chemical treatments. (**a**) Molecular model illustrating GAC covalently attached to peptidoglycan via a phosphodiester bond. GAC contains a →3)α-Rha(1→2)α-Rha(1→ repeating backbone. The β-GlcNAc side-chains are linked to the 3 position of the α-1,2-linked Rha ^95,96^. Phosphate groups in GAC are involved in the phosphodiester bond linking glycerol to the GlcNAc side-chain and the GAC reducing terminal sugar residue to peptidoglycan ^15^. (**b**) Release of GAC from GAS cell wall by mild acid hydrolysis or HONO deamination, before and after chemical *N*-acetylation. The treatment conditions are indicated by grey rectangles below the bar graph. (**c**) Release of GAC from GAS sacculi by mild acid hydrolysis, before (left) and after (right) chemical *N*-acetylation. (**d**) Release of GAC from GAS sacculi by HONO deamination, before (left) and after (right) chemical *N*-acetylation. In **b, c** and **d**, the concentration of GAC released from the sacculi was estimated by the modified anthrone assay described in Methods and normalized to total GAC content in sacculi. Symbols and error bars represent the mean and S.D. respectively (n=3 biologically independent replicates). Data are mean values ± S.D., n = 3 biologically independent experiments. In **b**, *P* values were calculated by one-way ANOVA with Tukey’s multiple comparisons test. In **c** and **d**, *P* values were calculated by two-way ANOVA with Tukey’s multiple comparisons test.

This proposed assembly mechanism is consistent with other isoprenol-mediated polysaccharide biosynthetic pathways in Gram-positive bacteria. The enzymes involved in polyrhamnose biosynthesis and GroP attachment are conserved and organized in similar gene clusters in many streptococci suggesting significant similarities in the structures of streptococcal rhamnopolysaccharides ^1^. However, several details of this pathway are still speculative and require further research. Importantly, the chemical structure of the linkage of GAC to peptidoglycan has not been addressed. In this study, in the course of developing a facile method for the efficient production of undegraded, highly purified GAC, suitable for use as a synthetic immunogen, we discovered that GAC was attached to peptidoglycan via an unexpectedly stable linkage unit. We found that this anomalous behavior is due to the presence of a substantial portion of N-unsubstituted glucosamine (GlcN) in the GAC-peptidoglycan linkage region. Using genetic, biochemical, and structural approaches, we identified PplD, encoded by *spy_0818* in the GAS 5005 genome, as the enzyme responsible for deacetylation of GlcNAc in the GAC linker unit. Furthermore, we revealed that deacetylation of the cell wall GlcNAc residues protects GAS from host small cationic antimicrobial proteins (AMPs).

## Results

### The GAC-peptidoglycan linkage is resistant to mild acid but sensitive to nitrous acid deamination

GAC was presumed to be covalently linked to peptidoglycan via a GlcNAc phosphodiester to the 6-hydroxyl group of *N*-acetylmuramic acid (MurNAc). This hypothesis is based on the observation that the biosynthetic pathways of GAC, and the *S. aureus* and *B. cereus* WTAs share initiating steps catalyzed by GacO homologs, providing the first sugar residue GlcNAc for the polysaccharide biosynthesis (Supplementary Fig. 1) ^2,8,13,17^. In the final steps of peptidoglycan decoration with WTAs, the LCP phosphotransferases attach GlcNAc 1-phosphate to peptidoglycan MurNAc residues forming a GlcNAc phosphodiester bond to peptidoglycan ^18^. The WTA-peptidoglycan phosphodiester bond is acid labile ^19^, and this aspect of the *S. aureus* and *B. subtilis* cell walls has been used to prepare undegraded, highly-enriched WTAs, freed of peptidoglycan remnants ^20,21^. Accordingly, the GAC-peptidoglycan linkage should also be sensitive to these conditions. We subjected purified GAS cell wall and intact sacculi to mild acid conditions (0.02 N HCl, 100ºC, 20 min), known to hydrolyze GlcNAc 1-phosphate, and measured the release of water-soluble GAC. Surprisingly, these well-established conditions released less than 25 % of GAC from GAS (Fig. 1 **b, c**). The resistance of GAC to release by mild acid hydrolysis suggested the presence of an un-acetylated amino sugar in the linkage region as has been reported for *Lactobacillus plantarum*, which contains a GlcN 1-phosphate-linked WTA. A phosphodiester bond connecting the *L. plantarum* WTA to peptidoglycan is resistant to mild acid hydrolysis unless first chemically *N*-acetylated ^20^. The chemical basis for the extraordinary resistance of GlcN 1-phosphate to acid hydrolysis is illustrated in Supplementary Fig. 2 **a, b**. Hydrolysis of the glycosidic bond of GlcNAc 1-phosphate is facilitated by protonation of the O1-oxygen and subsequent formation of a resonance stabilized oxonium ion intermediate (Supplementary Fig. 2 **a**). However, this reaction is dramatically slowed during incubation of GlcN 1-phosphate under these conditions, because the positively charged GlcN-ammonium group destabilizes the adjacent oxonium ion intermediate (Supplementary Fig. 2 **b**). To test the possibility that a hexosamine residue, devoid of an *N*-acetyl group, is present in the GAC-peptidoglycan linkage, purified GAS cell wall or sacculi were first chemically *N*-acetylated and then subjected to mild acid hydrolysis. GAC was nearly quantitatively released from peptidoglycan under these conditions, supporting our hypothesis (Fig. 1 **b, c**).

Sugars with primary amino groups, lacking N-acetyl groups, undergo deamination by reaction with nitrous acid (HONO). Deamination of polysaccharides containing GlcN residues results in the formation of 2,5-anhydromannose and hydrolysis of the glycosidic bond ^22^, as illustrated in Supplementary Fig. 2 **c**. This selective cleavage has been utilized successfully to release the *L. plantarum* WTA from peptidoglycan ^20^. Nitrite is rapidly protonated to form HONO in acidic conditions ^23^. We observed that incubation of isolated cell wall material and intact sacculi with acidified nitrite released 70-80% of GAC in a water-soluble form (Fig. 1 **b, d**). In contrast, chemically *N*-acetylated GAS cell wall or sacculi were resistant to this treatment (Fig. 1 **b, d**). Taken together, these results strongly suggest that GAC is attached to peptidoglycan via a hexosamine 1-phosphate linkage.

### The GAC-peptidoglycan linkage region contains GlcN and GlcNAc

Our observation that HONO deamination cleaves only 70-80% of GAC from peptidoglycan, and that treatment with mild acid released ∼20-25 % of GAC suggests that GAS produces two classes of rhamnopolysaccharide composed of major and minor fractions containing either hexosamine or *N*-acetylated hexosamine, respectively, in the linkage region. Importantly, mild acid hydrolysis of chemically *N*-acetylated cell wall released GAC nearly quantitatively (Fig. 1 **b**), indicating that failure to achieve complete release of GAC by these treatments was not due to incomplete hydrolysis of GAC from peptidoglycan or poor recovery of released material. To verify this conclusion, we sequentially subjected previously deaminated cell walls to mild acid hydrolysis and measured the release of soluble GAC (Supplementary Fig. 3). We observed that the HONO-resistant rhamnopolysaccharide fraction was fully released by the subsequent mild acid treatment. Furthermore, deamination of previously mild acid treated cell walls cleaves the remaining ∼80 % of the GAC which is resistant to mild acid hydrolysis (Supplementary Fig. 3).

To identify the hexosamine at the reducing end of the acetylated and un-acetylated forms of GAC, the polysaccharide was released from GAS cell wall by either deamination with HONO (nitrous acid sensitive GAC, or GAC^NA^) or mild acid hydrolysis (mild acid sensitive GAC, or GAC^MA^), and reduced chemically with sodium borohydride. Sodium borohydride reduces the aldehyde of 2,5-anhydromannose to form 2,5-anhydromannitol and the hemiacetal of GlcNAc to form *N*-acetyl-D-glucosaminitol (GlcNAcitol). The reduced GAC^NA^ and GAC^MA^ preparations were then fractionated by size-exclusion chromatography (SEC) on BioGel P150. As confirmed by the composition analysis, both treatments released similar, uniformly-sized polysaccharides eluting near the middle of the column elution volume (Fig. 2 **a** and **b**). When column fractions were assayed by GC-MS as TMS-methyl glycosides, the GAC^NA^ and GAC^MA^ preparations were found to contain Rha and GlcNAc as the only component sugars detected (Fig. 2 **a** and **b**, upper panels). The reducing-end sugars were detected as alditol acetates by GC-MS analysis. This method provides better detection of these minor terminal sugars in the presence of high concentrations of the polymer sugars, Rha and GlcNAc. We identified 2,5-anhydromannitol in the GAC^NA^ preparations, and GlcNAcitol in the GAC^MA^ preparations (Fig. 2 **a** and **b**, bottom panels), co-eluting with the peak of carbohydrate, as the only sugar alcohols in these fractions, strongly suggesting that the extracted GAC material contains GlcN and GlcNAc residues as the reducing moiety. Notably, we observed a lower apparent recovery of 2,5-anhydromannitol in the HONO-released fraction (Fig. 2 **b**), compared to the recovery of GlcNAcitol in the mild acid released material (Fig. 2 **a**). This result is expected because 2,5-anhydromannitol is not the sole product of deamination of hexosamines ^22,24^. In agreement, when we treated commercially available GlcN 1-phosphate with HONO, only ∼ 25% of the expected 2,5-anhydromannitol was recovered (Supplementary Figure 4).

**Fig. 2.**
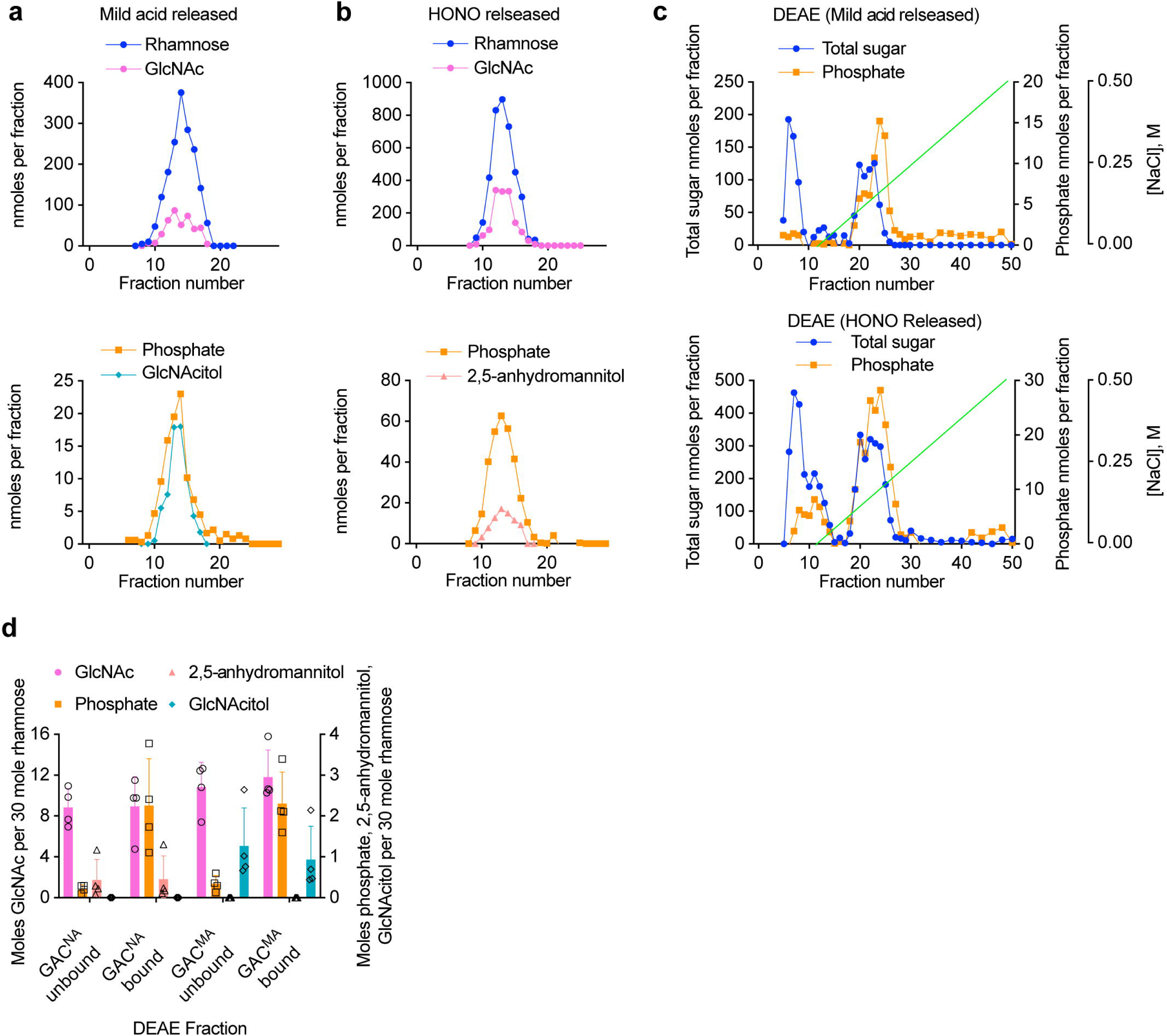
Glycosyl composition analysis of GAC released from cell wall by the chemical treatments and purified by size-exclusion and ion-exchange chromatography. (**a**) Size-exclusion chromatography of GAC released from GAS cell wall by mild acid hydrolysis or (**b**) deamination with HONO. Upper and lower panels show the composition of the BioGel P150 fractions of the individual GAC preparations. Prior to the chromatography, the extracted GAC material was reduced chemically with sodium borohydride. Fractions were analyzed for phosphate content by malachite green assay following digestion with perchloric acid. Rha and GlcNAc contents were measured by GC-MS as TMS-methyl-glycosides. GlcNAcitol and 2,5 anhydromannitol contents were estimated by GC-MS as alditol acetates. The chromatographic profiles are representative of more than three separate experiments. (**c**) Ion-exchange chromatography of GAC released from the GAS cell wall by mild acid (upper panel) and HONO (lower panel). Fractions containing the GAC material from Fig. 2 **a** and **b** were pooled, concentrated, desalted by spin column, loaded onto DEAE-Toyopearl and eluted with a NaCl gradient (0-0.5 M). Fractions were analyzed for total sugar content by anthrone assay and total phosphate content by malachite green assay following perchloric acid digestion. The experiments were performed at least three times and yielded the same results. Data from one representative experiment are shown. (**d**) Glycosyl composition analysis of the GAC material purified by ion-exchange chromatography. The GAC material released by either deamination with HONO (GAC^NA^) or mild acid hydrolysis (GAC^MA^) was analysed as shown in Fig. 2 **c**. Fractions unbound (flow-through) and bound (eluted with a NaCl gradient) to the DEAE column were pooled, concentrated, desalted by spin column and analyzed by GC-MS to determine the Rha, GlcNAc, 2,5-anhydromannitol and GlcNAcitol concentrations as described above. Malachite green assay was used to assay the phosphate concentration. The concentrations of GlcNAc, 2,5-anhydromannitol, GlcNAcitol and phosphate are expressed as moles per 30 moles of Rha. Columns and error bars represent the mean and S.D., respectively (n = 3 biologically independent replicates).

We have previously shown that GAS and *Streptococcus mutans* produce rhamnopolysaccharides with variable GroP content, and these forms can be separated by ion-exchange chromatography ^15,25^. Further experiments were performed to examine whether the GlcN and GlcNAc linker residues are restricted to a specific ionic isoform of GAC. Fractionation of GAC^NA^ and GAC^MA^ on DEAE Toyopearl revealed that both polysaccharides contained a similar array of ionic forms: a neutral glycan, not bound to the ion-exchange column, and a strongly bound component (Fig. 2 **c**). Similar heterogeneity of GAC was previously observed when the polysaccharide was cleaved from peptidoglycan by the peptidoglycan hydrolases ^15^. As expected, the neutral glycan contains very low phosphate content indicating that it has a low degree of GroP modification (Fig. 2 **c** and **d**). The glycosyl composition of the pooled GAC fractions revealed that the column-bound and unbound fractions from both polysaccharide preparations contain similar Rha/GlcNAc ratio (Fig. 2 **d**). Furthermore, 2,5-anhydromannitol was detected in both the column-bound and unbound fractions of the GAC^NA^ preparations, and GlcNAcitol in both the column-bound and unbound fractions of the GAC^MA^ preparations (Fig. 2 **d**). These observations provide evidence that the GlcN and GlcNAc linker isoforms are not specifically linked to the presence or absence of GroP in GAC and that the only apparent difference in the two populations of GAC is the acetylation status of the linkage GlcN unit.

### Chemical structure of the GAC-peptidoglycan linkage region

To elucidate the exact chemical structure of the linkage region, GAC liberated from peptidoglycan by the peptidoglycan hydrolases, PlyC and mutanolysin ^15^, was subjected to NMR analysis. As expected from our previous study ^15^, when GAC material was analyzed by ^1^H,^13^C-HSQC NMR, the GAC trisaccharide repeating units, →3)-α-L-Rha*p*-(1→2)[β-D-Glc*p*NAc-(1→3)]-α-L-Rha*p*-(1→, were observed at highest level of intensity. The polysaccharide decorated by *sn*-Gro-1-phosphate groups as substituents at O6 of the side-chain D-Glc*p*NAc residues were identified at the second intensity level of NMR spectra ^15^. To unravel how GAC is linked to peptidoglycan, the analysis was performed at the third level of intensity in the NMR spectra. In the ^31^P NMR spectrum of GAC, a resonance was observed at −1.35 ppm, having a chemical shift characteristic of a phosphodiester ^26^. From ^1^H,^31^P-Hetero-TOCSY experiments ^27^ correlations may be detected from the ^31^P nucleus to protons of adjacent residues and in the spectrum with a mixing time of 50 ms cross-peaks were observed to protons of a methylene group, δ_H_ 4.10 and 4.19, identified in a multiplicity-edited ^1^H,^13^C-HSQC spectrum (δ_C6_ 64.3) (Fig. 3); these correlations were confirmed via ^1^H,^31^P-HMBC experiments. A longer mixing time of 80 ms in the ^1^H,^31^P-HMBC experiment and analysis of ^1^H,^1^H-TOCSY spectra for which mixing times up to 120 were used in conjunction with ^1^H,^13^C-HSQC, ^1^H,^13^C-HSQC-TOCSY and 2D ^1^H,^1^H-NOESYspectra, complemented by NMR chemical shift predictions made by the CASPER program, ^28^ facilitated identification of a hexopyranoside (δ_C1_ 102.4) spin-system having the *gluco*-configuration. Importantly, in the ^1^H,^13^C-HMBC spectrum a correlation was observed between δ_Hα_ 4.36 of a lactyl group and δ_C3_ 80.2 of the hexopyranoside residue and in a ^1^H,^1^H-NOESY spectrum connectivity between entities was further substantiated by a cross-peak at δ_H__α_ 4.36/δ_H3_ 3.63 thereby identifying the sugar having δ_C4_ 75.7 and ^1^*J*_C1,H1_ 167 Hz as a β-MurNAc residue being part of a peptidoglycan chain (Supplementary Table 1). The ^13^C NMR resonance assigned to C5 of the MurNAc residue showed ^3^*J*_C5,P_ 7.6 Hz, fully consistent with a phosphodiester 6-*O*-substitution ^29^. The spectral data of nuclei in the hexopyranose ring are consistent with previously reported NMR chemical shift assignments of an oligomeric form of peptidoglycan structures ^30^.

**Fig. 3.**
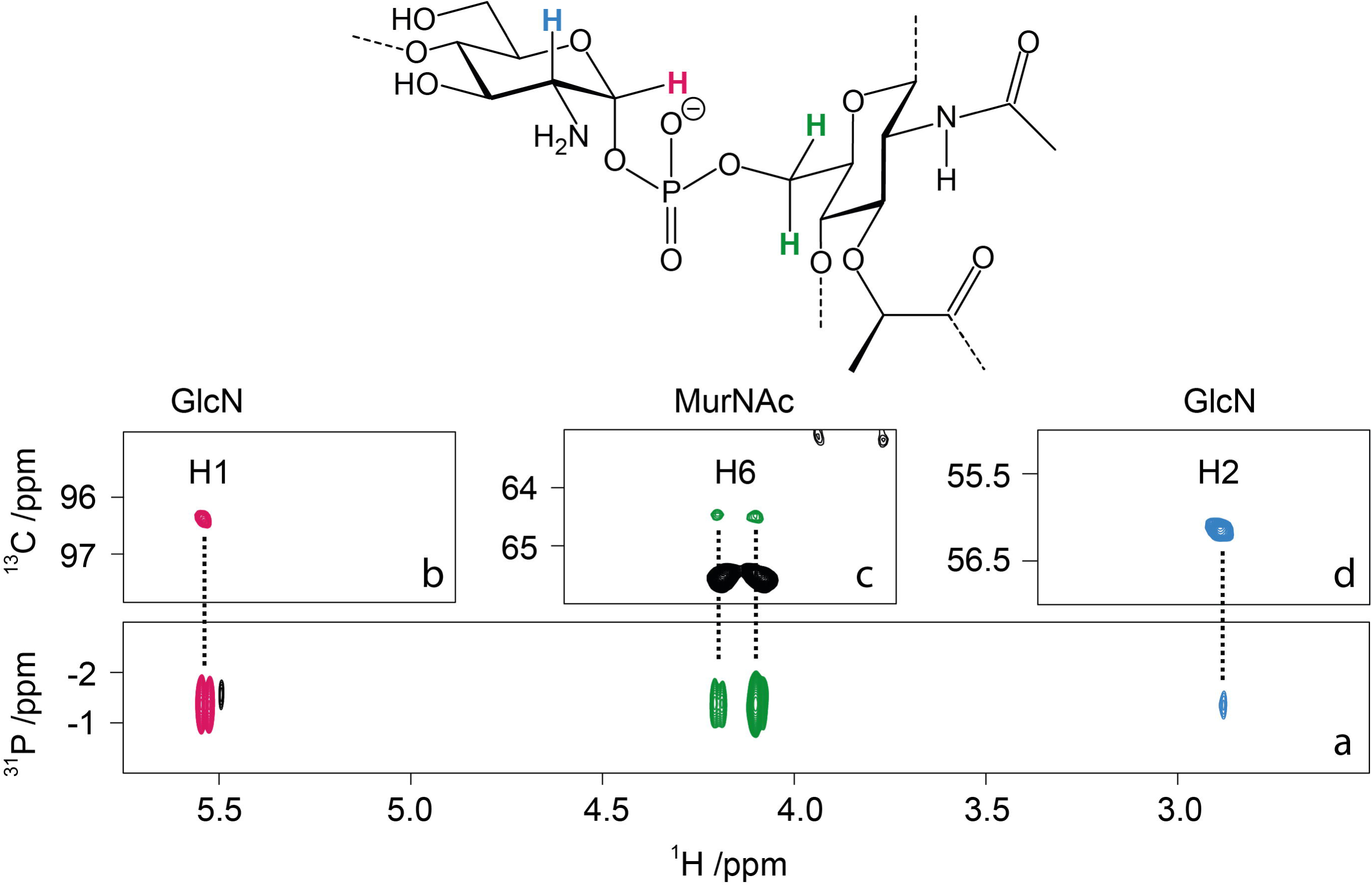
Chemical structure of the GAC-peptidoglycan linkage region determined by NMR analysis. Representative structure of the linkage region of GAC anchored to the peptidoglycan layer (**top**). ^1^H,^31^P-HMBC NMR spectrum (**a**) with a 90 ms delay for the evolution of long-range *J* couplings shows the key correlation between proton H1 (**b**) and H2 (**d**) of GlcN, and the two protons in position 6 of MurNAc (**c**) to a unique ^31^P signal at −1.35 ppm, characteristic of a phosphodiester linkage between the two residues. Panels (**b**), (**c**) and (**d**) show the corresponding signals in the multiplicity-edited ^1^H,^13^C-HSQC NMR spectrum. The two H6 signals of MurNAc were observed in opposite phase due to the multiplicity selection.

The ^1^H,^31^P-Hetero-TOCSY spectra employing a mixing time of 50 ms also showed correlations from ^31^P to a proton at δ_H1_ 5.54 and with a longer mixing time of 80 ms a cross-peak was identified at δ_H2_ 2.89 (Fig. 3); these correlations were confirmed in a ^1^H,^31^P-HMBC spectrum. The combined analysis of ^1^H,^1^H-TOCSY and ^1^H,^13^C-HSQC spectra indicated that the spin-system originates from a hexopyranoside residue (δ_C1_ 96.5) having the *gluco*-configuration with an α-anomeric configuration (^1^*J*_C1,H1_ 177 Hz) as part of a phosphodiester linkage ^31,32^. The ^13^C NMR chemical shift δ_C2_ 56.0 is indicative of a nitrogen-carrying carbon suggesting that this residue is the primer of the GAC ^16^. However, the ^1^H NMR chemical shift δ_H2_ 2.89 at pD ∼8 is conspicuously low and was pD-dependent, which supports that the D-GlcN residue is devoid of an *N*-acetyl group ^33-35^. Furthermore, a β-L-Rha*p* residue was identified by ^1^*J*_C1,H1_ 163 Hz and intra-residue cross-peaks in a ^1^H,^1^H-NOESY spectrum from H1 to H2, H3 and H5. The ^13^C NMR chemical shift of C3 at δ_C3_ 81.4 in the β-L-Rha*p* residue (Supplementary Table 1) is higher by 7.6 ppm compared to that of the monosaccharide ^36^, suggesting that it is substituted by a sugar residue at O3 ^37^. The priming D-GlcN residue is substituted at O4 by the β-L-Rha*p* residue as seen from a ^1^H,^13^C-HMBC correlation δ_H1_ 4.89/δ_C4_ 77.7 and in a ^1^H,^1^H-NOESY spectrum with a cross-peak at δ_H1_ 4.89/δ_H4_ 3.69, consistent with a β-L-Rha*p*-(1→4)-α-D-Glc*p*N disaccharide, derived from transfer of L-Rha from TDP-β-L-Rha onto the acceptor GlcNAc-PP-Und catalyzed by GacB ^16^ (Supplementary Fig. 1). Taken together, the results from the NMR analysis show that GAC is attached by its primer D-GlcN via a phosphodiester linkage to MurNAc of the peptidoglycan.

### The newly synthesized cell wall is decorated with GlcNAc 1-phosphate-linked GAC

In Gram-positive bacteria, the cell wall polysaccharides conceal peptidoglycan from molecular interactions with proteins ^25,38^. We reasoned that since the chemical treatments of GAS sacculi cleave a specific form of GAC possessing either the GlcN or GlcNAc residues in the GAC-peptidoglycan linkage, these treatments might reveal surface peptidoglycan regions expressing the pertinent modification by following peptidoglycan-specific protein binding. Thus, to examine the cellular localization of these specific GAC forms, we generated a fluorescent molecular tool with tight binding to peptidoglycan. We constructed a protein fusion, GFP-AtlA^Efs^, in which the C-terminal cell wall-binding domain of the *Enterococcus faecalis* major cell division autolysin AtlA ^39,40^ is fused at the N-terminus with a green fluorescent protein (GFP) (Supplementary Figure 5). This cell wall-binding domain consists of six LysM repeats (Pfam PF01476) that recognize peptidoglycan ^41^. The protein fusion was added exogenously to the GAS sacculi subjected to either no treatment, mild acid hydrolysis, or deamination. Fluorescence microscopy imaging revealed that the protein fusion did not associate with the majority of untreated sacculi (Fig. 4 and Supplementary Table 2). In contrast, the sacculi treated with HONO demonstrated uniform distribution of GFP-AtlA^Efs^ over the cell surface (Fig. 4). Interestingly, the majority of sacculi subjected to mild acid hydrolysis were stained by GFP-AtlA^Efs^ in the regions of active cell division: septal regions and newly formed poles (Fig. 4 and Supplementary Table 2). This observation indicates that i) the chemical treatments do not lead to degradation of the peptidoglycan sacculi; and ii) the regions of active cell division contain the GlcNAc 1-phosphate-linked GAC. These zones correspond to the newly synthesized cell wall.

**Fig. 4.**
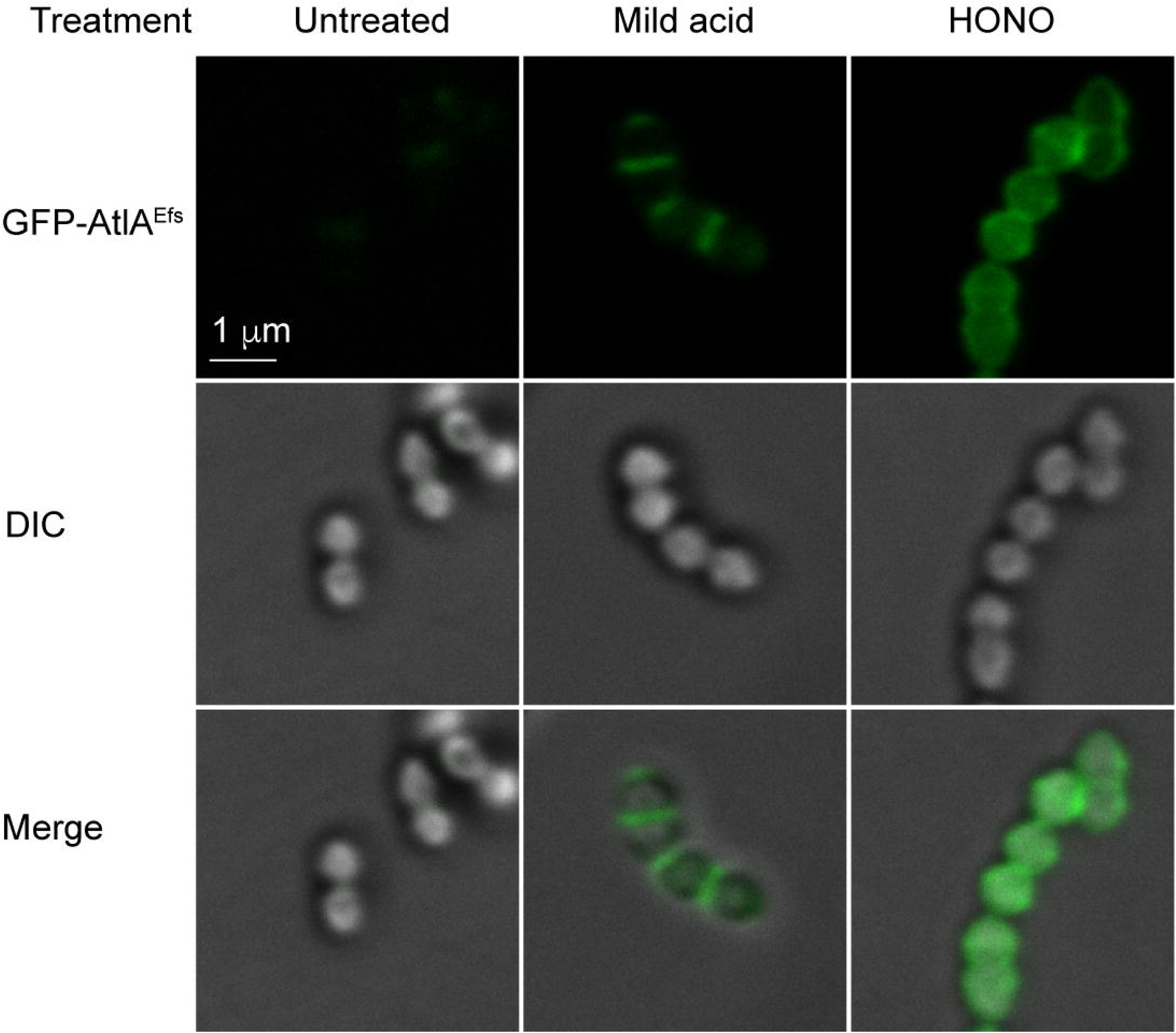
Binding of GFP-AtlA^Efs^ to GAS sacculi. Intact sacculi (untreated), sacculi subjected to mild acid hydrolysis (mild acid), or deamination with HONO (HONO) were incubated with GFP-AtlA^Efs^, and examined by fluorescence microscopy (top panels) and DIC (middle panels). An overlay of fluorescence and DIC is shown in the bottom panels. The experiments were performed independently three times and yielded the same results. Representative image from one experiment is shown. Scale bar is 1 μm.

### PplD deacetylates the GAC-peptidoglycan linkage region

Our observation that a substantial fraction of the GAC-peptidoglycan linkage region carries a GlcN residue suggests enzymatic deacetylation of GlcNAc as an undescribed step in GAC biosynthesis. The GAS genome encodes two proteins, PgdA and Pdi, annotated as putative GlcNAc deacetylases. PgdA is a homolog of the peptidoglycan GlcNAc N-deacetylase from *Streptococcus pneumoniae* ^42^. Pdi was originally identified in the fish pathogen *Streptococcus iniae* ^43^. While the function of this protein has not been described, it was named Pdi for <u>p</u>olysaccharide <u>d</u>eacetylase of *S. <u>i</u>niae*. We renamed this protein to *PplD* (<u>p</u>olysaccharide-<u>p</u>eptidoglycan <u>l</u>inkage <u>d</u>eacetylase) for the reasons provided below. To examine the functions of PplD and PgdA in deacetylation of the GAC linker region, we generated *pplD* and *pgdA* single deletion mutants and the double deletion mutant in a GAS NZ131 strain, creating GASΔ*pplD*, GASΔ*pgdA* and GASΔ*pplD*Δ*pgdA*, respectively. Since the chemical treatments produced similar results with the GAS cell wall and sacculi (Fig. 1 **b, c** and **d**), for convenience we used the sacculi of the constructed strains to test the sensitivity of the phosphodiester linkage to mild acid and HONO treatment. Mild acid hydrolysis liberated the polysaccharide almost quantitatively from the GASΔ*pplD* and GASΔ*pplD*Δ*pgdA* sacculi (Fig. 1 **c**). Conversely, the sacculi of these mutants were almost completely resistant to deaminative cleavage with HONO (Fig. 1 **d**). Moreover, chemical *N*-acetylation of the GASΔ*pplD* and GASΔ*pplD*Δ*pgdA* sacculi prior to the treatments with either mild acid or HONO did not significantly affect the release of GAC (Fig. 1 **c** and **d**). In contrast, the GASΔ*pgdA* sacculi behaved in these assays like the WT control (Fig. 1 **c** and **d**). These data indicate that PplD is required for deacetylation of the GAC-peptidoglycan linker region.

### PplD structure

PplD is predicted to contain an N-terminal transmembrane helix and an extracellular catalytic domain (ePplD) connected by a linker region. To facilitate the functional analysis of PplD, we solved the crystal structure of ePpLD (residues 94-320) to 1.78 Å resolution (Supplementary Table 3). There were two molecules in the asymmetric unit that are very similar to each other with a r.m.s.d. of 0.3 Å over 226 aligned Cα atoms. Analysis of the crystal contacts revealed that the dimer interface is the largest with 632 Å^2^ buried surface area that scored below the stable complex formation criteria ^44^. In addition, ePplD behaves as a monomer in solution according to SEC. The structure of ePplD revealed a (β/α)_7_-barrel fold that is characteristic of carbohydrate esterases of the CE4 family (Fig. 5 **a**). The analysis of the putative active site revealed the presence of a metal ion, which we identified as a Zn^2+^ based on analysis of anomalous difference maps and temperature factors of the ion and coordinating atoms. The Zn^2+^ ion is coordinated by the side chains of residues D168, H223 and H227 (Fig. 5 **b**). This coordination pattern of the D-H-H residues is common for the metal coordination in CE4 family members. An additional electron density observed next to the Zn^2+^ ion was assigned as acetate, one of the products of the deacetylation reaction. The compound makes a bi-dentate contact with the Zn^2+^ ion. Altogether, the side chains of D168, H223 and H227, acetate and a water molecule complete an octahedral coordination of the Zn^2+^ ion (Fig. 5 **b**). One of the oxygen atoms of acetate makes a hydrogen bond with the backbone amide of S264, an oxyanion hole, while another oxygen atom of acetate forms a hydrogen bond with catalytic residue H105. A semi-buried D167, which is likely involved in coordination of a catalytic water molecule, makes a bi-dentate salt bridge with R302 and an additional hydrogen bond with N191. Altogether, the structural features of ePplD are consistent with the deacetylase function of this enzyme.

**Fig. 5.**
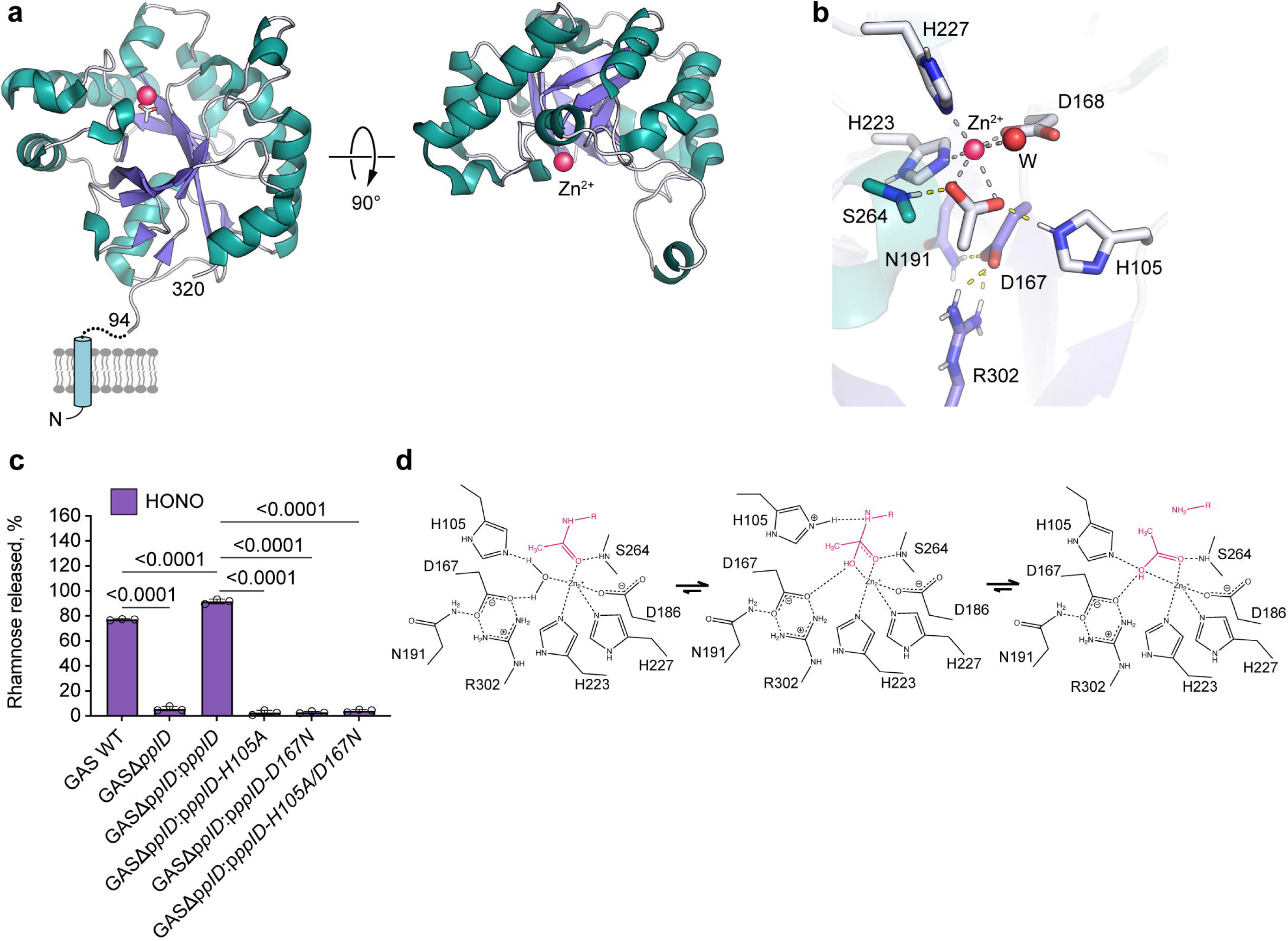
PplD deacetylates GlcNAc in the GAC-peptidoglycan linkage region. (**a**) Predicted topology of PplD showing a transmembrane helix and structure of extracellular domain with the enzymatic active site is depicted on left panel. Right panel depicts the structure of ePplD viewing at the active site with the Zn^2+^ ion shown as a magenta sphere. (**b**) A close-up view of the active site PplD structure in complex with acetate. (**c**) The sacculi purified from GAS WT, GASΔ*pplD*, GASΔ*pplD*:p*pplD*, GASΔ*pplD*:p*pplD-H105A*, GASΔ*pplD*:p*pplD-D167N* and GASΔ*pplD*:p*pplD-H105A/D167N* were subjected to deamination with HONO. The concentration of GAC released from the sacculi was estimated by the modified anthrone assay and normalized to total GAC content in the analyzed materials. Symbols and error bars represent the mean and S.D., respectively (n = 3 biologically independent replicates). *P* values were calculated by one-way ANOVA with Tukey’s multiple comparisons test. (**d**) Proposed catalytic mechanism for PplD-mediated deacetylation of the GAC-peptidoglycan linkage region.

The structural comparison of PplD with the available protein structures shows that this enzyme is more distantly related to *S. pneumoniae* PgdA ^45^, but similar to the exopolysaccharide poly-*N*-acetylglucosamine (PNAG) deacetylases PgaB and IcaB produced by *Escherichia coli* and *Ammonifex degensii*, respectively, (Supplementary Table 4) ^46,47^. The core of the catalytic domains of PplD and PgdA is similar with a r.m.s.d. of 2.4 Å over 140 Cα atoms. However, the N-and C-termini are swapped and located on the opposite ends of β/α barrel (Supplementary Fig. 6). Moreover, in comparison to canonical members of the CE4 protein family, PplD displays differences in the MT1-MT5 sequence motifs. While, the MT1, MT2 and MT5 motifs containing the catalytic residues and the residues coordinating Zn^2+^ ion are conserved in PplD and PgdA (Supplementary Fig. 6), PplD lacks the MT4 motif that carries an aspartic acid residue involved in activation of the catalytic histidine (H105 in PplD). Additionally, the MT3 motif of PplD lacks an arginine residue that coordinates a catalytic aspartic acid residue (D167 in PplD). Instead, D167 is coordinated by R302 located at a different position in the sequence. These sequence deviations from canonical CE4 hydrolases are also present in *E. coli* PgaB and *A. degensii* IcaB.

To functionally assess the requirement of the presumed catalytic residues of PplD, we complemented GASΔ*pplD* and GASΔ*pplD*Δ*pgdA* with WT *pplD* expressed on a plasmid creating the GASΔ*pplD*:p*pplD* and GASΔ*pplD*Δ*pgdA*:p*pplD* strains, and GASΔ*pplD* with catalytically inactive versions of *pplD* in which the active site residues H105 and D167, were replaced by alanine and asparagine, respectively, creating GASΔ*pplD*:p*pplD-H105A*, GASΔ*pplD*:p*pplD-D167N* and GASΔ*pplD*:p*pplD-H105A/D167N*. Complementation of the mutants by expression of the WT *pplD* completely restored the phenotype of the mutants (Fig. 1 **c, d** and 5 **c**). Importantly, in GASΔ*pplD*:p*pplD* and GASΔ*pplD*Δ*pgdA*:p*pplD*, the phosphodiester GAC-peptidoglycan bond was significantly more sensitive to deaminative cleavage by HONO than in the WT (Fig. 1 **d** and 5 **c**). This result indicates that the plasmid expression of PplD in GAS provided the cells with fully deacetylated GAC linker region. In contrast, the GASΔ*pplD*:p*pplD-H105A*, GASΔ*pplD*:p*pplD-D167N* and GASΔ*pplD*:p*pplD-H105A/D167N* mutants behaved similarly to GASΔ*pplD* (Fig. 5 **c**). Thus, based on the functional requirement of H105 and D167 for catalysis and the structural features of the active site, we propose that PplD functions via an alternative enzymatic mechanism similar to *A. degensii* IcaB ^47^. In this mechanism, the hydrolysis of the GlcNAc moiety would occur via a water molecule coordinated by Zn^2+^ ion and D167 (Fig. 5 **d**). The nucleophilic attack on the *N*-acetyl group of the GlcNAc residue would be initiated by deprotonation of the catalytic water molecule by H105. The resulting tetrahedral oxyanion intermediate would be coordinated by the Zn^2+^ ion and the backbone amide of S264 acting as an oxyanion hole. The protonation of the nitrogen of the intermediate by H105 in the histidinium form would lead to hydrolysis and generation of the GlcN moiety and a free acetate. In this mechanism, H105 would serve as a bifunctional catalytic base/acid of the reaction.

### Modulation of the cell wall net charge by PplD, PgdA and GacH affects resistance to AMPs

In a number of bacteria, PgdA protects bacteria from host AMP, lysozyme, by deacetylating the peptidoglycan GlcNAc residues ^42,48-50^. Lysozyme catalyzes the breakdown of β-(1,4) linkages between the MurNAc and GlcNAc saccharides leading to peptidoglycan hydrolysis and bacterial lysis ^51^. In addition to *N*-acetylmuramidase activity, lysozyme kills bacteria by a non-enzymatic mechanism ^52^. This mechanism is similar to killing by small cationic peptides ^53^, and is attributed to lysozyme crossing the cell wall and binding to negatively charged bacterial membranes thus leading to membrane permeabilization. Deacetylation of GlcNAc in peptidoglycan by PgdA directly alters lysozyme binding to its substrate negatively affecting lysozyme-mediated degradation of peptidoglycan. Interestingly, in *S. iniae*, the PplD homolog also confers resistance to lysozyme ^43^. Hence, to understand the role of cell wall deacetylation in protection of GAS against lysozyme, first, we assessed muramidase-dependent lysis of the WT GAS, GASΔ*pplD*, GASΔ*pgdA* and GASΔ*pplD*Δ*pgdA* cells treated with lysozyme. When GAS strains were incubated with 1 mg/ml lysozyme for 18 hours, no lysozyme-dependent decrease in turbidity was observed consistent with the established high resistance of GAS to lysozyme. In contrast, incubation of GAS strains with an alternative highly efficient N-acetylmuramidase, mutanolysin, produced a substantial drop in turbidity compared to control (Supplementary Fig. 7). Since the sensitivity of GAS to lysozyme appeared to be below the limits of detection detected by turbidity analysis, cell wall degradation was estimated with a more sensitive assay using dot-blot analysis for the water-soluble form of GAC released from the bacteria during digestion (Supplementary Fig. 8 **a)**. Lysozyme produced a very weak solubilization of GAC from WT GAS and GASΔ*pplD* cells, and more pronounced release of GAC from GASΔ*pgdA* and GASΔ*pplD*Δ*pgdA* cells (Supplementary Fig. 8 **b**). Furthermore, lysozyme released GAC from GASΔ*pplD*Δ*pgdA* more efficiently, than from GASΔ*pgdA* (Supplementary Fig. 8 **b)**. Altogether, these data indicate that lytic degradation of the GAS cell wall by lysozyme is minor due to the presence of GlcN in peptidoglycan as the result of PgdA catalytic activity. However, when PgdA is absent, deacetylation of the GAC linker region by PplD can be detected as a factor conferring protection against lysozyme-mediated lysis.

We previously reported that in comparison to the parental strain, the GroP-deficient mutant of GAS, GASΔ*gacH*, is significantly more resistant to a highly cationic enzyme human group IIA secreted phospholipase A_2_ (hGIIA sPLA_2_ hereafter hGIIA) ^15^. This phenotype is likely due to either decreased binding of hGIIA to the cell surface or reduced permeability of the cell wall to hGIIA. Since deacetylation of GlcNAc residues increases the cell wall net positive charge, it might lead to neutralization of negative charges conferred by GroP moieties and as a result, affect the translocation of cationic AMPs across the cell wall. To understand how modulation of the cell wall net charge by deacetylation of GlcNAc residues alters susceptibility of GAS to cationic AMPs, we analyzed the resistance of WT GAS, GASΔ*pplD*, GASΔ*pgdA*, GASΔ*pplD*Δ*pgdA* to the antimicrobial action of hGIIA, lysozyme, or histone mixture (Fig. 6 **a, b, c** and Supplementary Fig. 8 **c**). We observed that deletion of either *pplD* or *pgdA*, or both genes, yielded strains with enhanced susceptibility to hGIIA (Fig. 6 **a**). In contrast, deletion of *pplD* did not affect susceptibility of bacteria to lysozyme (Fig. 6 **b** and Supplementary Fig. 8 **c**) or a histone mixture (Fig. 6 **c)**. However, sensitivity to killing by these AMPs was significantly increased in GASΔ*pgdA* and GASΔ*pplD*Δ*pgdA*, with a more pronounced effect observed in GASΔ*pplD*Δ*pgdA*. Moreover, the expression of WT PplD in GASΔ*pplD* and GASΔ*pplD*Δ*pgdA* significantly increased the resistance of the parental strains to the AMPs, especially for hGIIA (Fig. 6 **a, b, c** and Supplementary Fig. 8 **c**). This observation correlates with the conclusion that the level of deacetylation of the GAC-peptidoglycan linkage region in the complemented strains is higher than in the WT strain, and indicates that both PgdA and PplD enhance resistance of GAS to the tested AMPs.

**Fig. 6.**
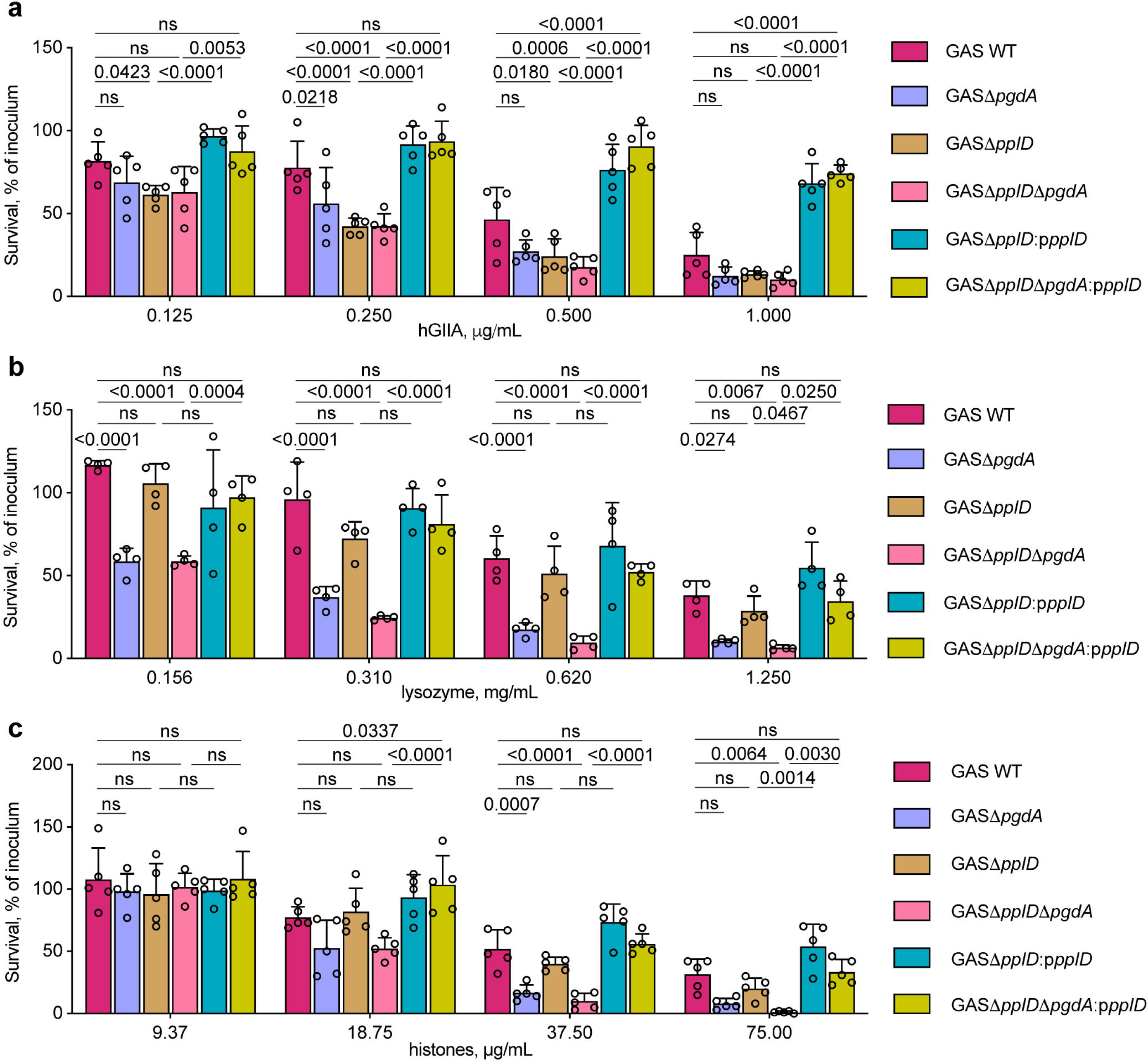
*N*-deacetylases PplD and PgdA contribute to protection of GAS against AMPs. (**a, b** and **c**) Analysis of resistance of GAC mutants deficient in *N*-deacetylation of GlcNAc mediated by PplD and PgdA to (**a**) hGIIA, (**b**) lysozyme, and (**c**) histone mixture. Data are mean values ± S.D., n = 5 biologically independent experiments in **a** and **c**, n = 4 biologically independent experiments in **b**. *P* values were calculated by two-way ANOVA with Bonferroni’s multiple comparison test.

Finally, we examined susceptibility of the GroP-deficient mutant, GASΔ*gacH*, and its complemented strain, GASΔ*gacH:*p*gacH*, to the antimicrobial action of lysozyme or histones to confirm that the mechanisms of lysozyme and histone killing of GAS are charge-dependent. Deletion of *gacH* rendered GAS more resistant to lysozyme and histones, and the phenotype was reversed by complementation of GASΔ*gacH* with WT *gacH* (Fig. 7 **a, b**). Collectively, our results are consistent with a model in which the addition of positive residues by deacetylation of GlcNAc in peptidoglycan and the GAC linkage region counteracts the cell wall net negative charge provided by GroP moieties resulting in reduced translocation of cationic AMPs across the cell wall (Fig. 7 **c**).

**Fig. 7.**
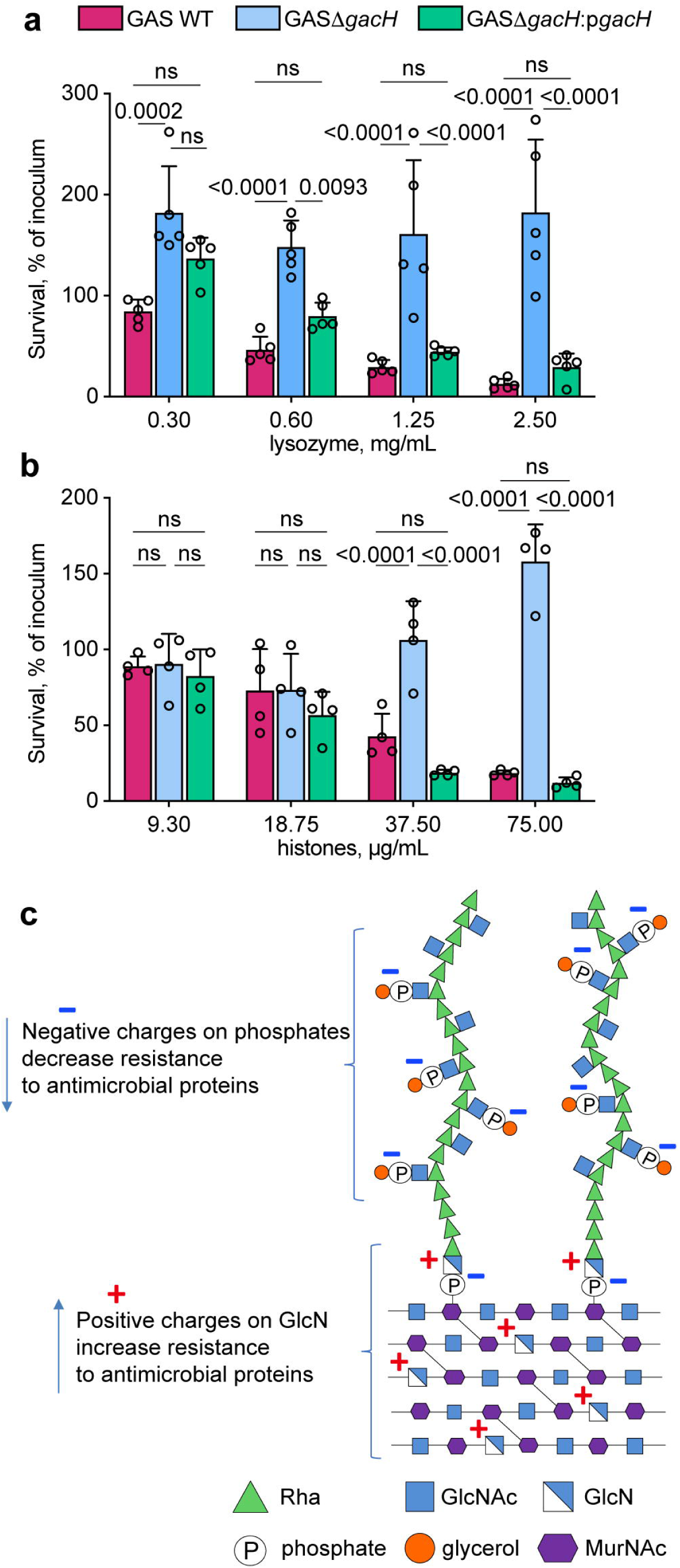
The role of the GroP modification in GAS protection against histones and lysozyme. (**a** and **b**) Analysis of resistance of GAC mutants deficient in the GroP modification to (**a**) lysozyme, and (**b**) histone mixture. Data are mean values ± S.D., n = 5 biologically independent experiments in **a**, n = 4 biologically independent experiments in **b**. *P* values were calculated by two-way ANOVA with Bonferroni’s multiple comparison test. (**c**) Introduction of charges to the GAS cell wall by GlcNAc deacetylation and GroP modification modulates resistance to host cationic antimicrobial proteins.

### PplD homologs deacetylate respective rhamnopolysaccharides in streptococci

To understand whether deacetylation of the polysaccharide-peptidoglycan linkage region is widespread in bacteria, we examined the distribution of the PplD family of proteins throughout the bacterial kingdom. PplD homologs were predominantly identified in the genomes of streptococcal species (Supplementary Fig. 9). Interestingly, members of the *S. mitis* group, that express choline-containing WTAs, and not Rha-containing polysaccharides, possess only the PgdA homolog, supporting the conclusion that PplD is only active on rhamnopolysaccharides. We also observed that genes encoding *pplD* homologs in a number of streptococcal strains contain a frame shift mutation and thus are likely non-functional. Hence, to investigate the function of PplD in streptococcal species, we selected for analysis the following strains: *Streptococcus agalactiae* (Group B Streptococcus, GBS) A909, *S. mutans* Xc, and *Streptococcus equi subsp. equi* CF32 in which *pplD* is an intact gene, and GBS COH1 and *Streptococcus thermophilus* LMG 18311 in which the *pplD* homologs have a frame shift mutation (Supplementary Fig. 10). Additionally, we deleted the *pplD* homologs in *S. mutans* Xc and GBS A909 strains creating the SMUΔ*pplD* and GBSΔ*pplD* strains. In agreement with the function of PplD in deacetylation of the polysaccharide-peptidoglycan linker, the rhamnopolysaccharides were liberated from the cell walls by HONO in the strains with the intact *pplD* (Fig. 8 **a, b)**. In contrast, this treatment did not cleave the polysaccharides from the strains carrying the inactivated *pplD*. However, these polysaccharides were released by mild acid hydrolysis (Fig. 8 **a, b)**. Furthermore, expression of the GAS WT PplD but not the catalytically inactive versions of PplD, PplD-D167N and PplD-H105A/D167N, restored the phenotype of SMUΔ*pplD* (Fig. 8 **a**). Taken together, the results support the conclusion that in the large group of streptococci expressing the Rha-containing polysaccharides, the polysaccharide-peptidoglycan linkage region is deacetylated by PplD homologs suggesting that PplD recognizes for catalysis the polyrhamnose linker region sugars that are conserved in many streptococci ^1,16^.

**Fig. 8.**
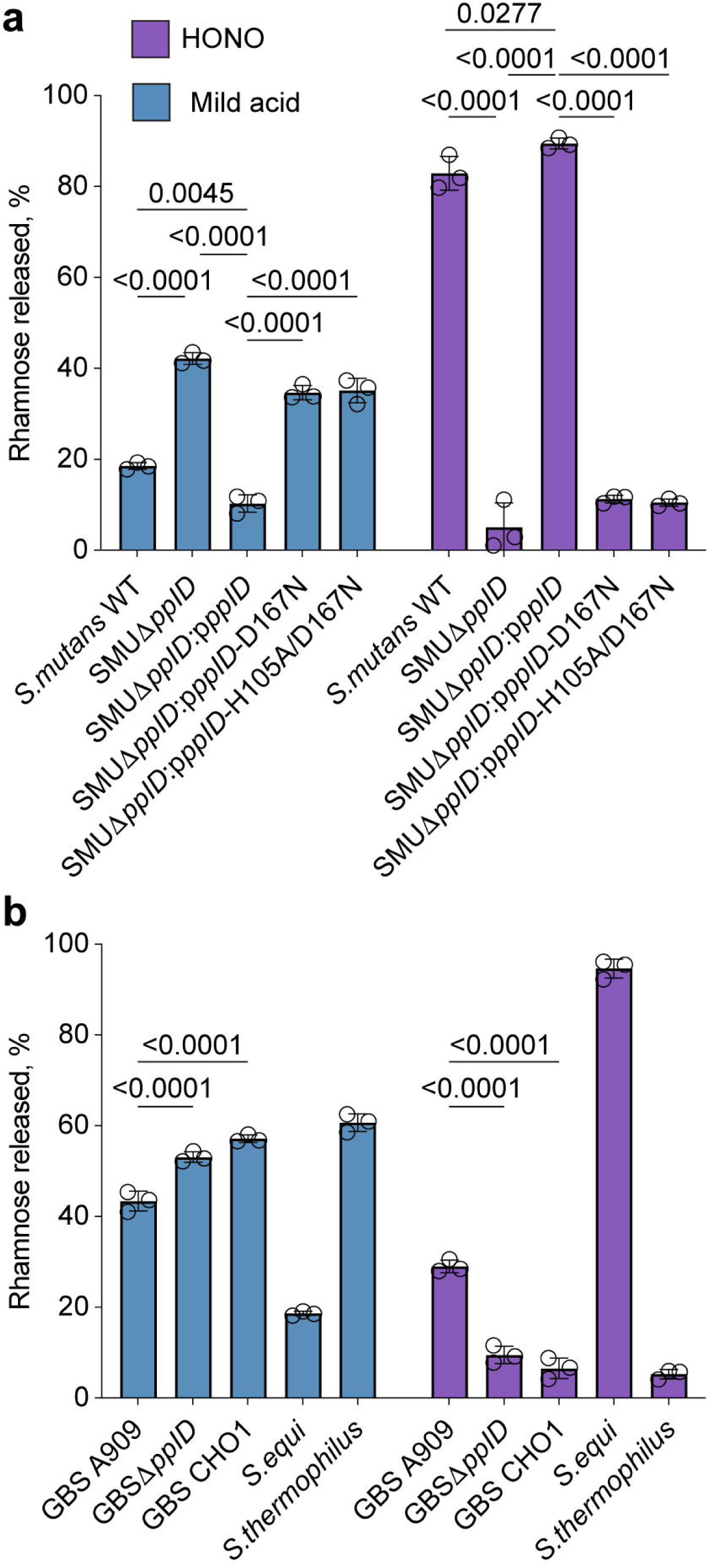
Analysis of PplD-mediated deacetylation in streptococcal species. (**a**) Release of SCC from the sacculi purified from *S. mutans* strains by mild acid hydrolysis (left) or deamination with HONO (right). (**b**) Release of the Rha-containing polysaccharides from the cell wall purified from GBS, *S. equi* and *S. thermophilus* strains by mild acid hydrolysis (left) or deamination with HONO (right). The concentration of polysaccharide released from the sacculi or cell wall was estimated by the modified anthrone assay and normalized to total content of polysaccharide in the starting material. Symbols and error bars represent the mean and S.D. respectively (n = 3 biologically independent replicates). *P* values were calculated by two-way ANOVA with Tukey’s multiple comparisons test.

## Discussion

Currently, no licensed vaccine exists for GAS which is a leading cause of morbidity and mortality worldwide. Glycoconjugate vaccines obtained by covalent linkage of bacterial polysaccharides to carrier proteins have been proven highly successful in the prevention of a number of infectious diseases ^54^. The major component of GAS cell wall, GAC, is an important determinant of viability and virulence, and is a promising target for the development of vaccines to treat GAS infections. One of the obstacles to the GAC-based vaccine development is the lack of efficient and structurally accurate methods of GAC conjugation to protein carriers ^9^, primarily as a result of misperceptions of the chemical nature of the GAC-peptidoglycan linkage unit. A consequence of this incomprehension is that previous methods for preparation of GAC have either been notably inefficient or relied on harsh chemical treatments for polysaccharide cleavage and the insertion of reactive chemical groups required for coupling reactions to carrier proteins ^6-11^. In this report, using NMR studies together with chemical degradation and glycosyl composition analysis, we demonstrate that a significant fraction of GAC contains GlcN 1-phosphate in the peptidoglycan linkage unit. This previously unrecognized structural feature of the GAC introduces a positive charge at the site of attachment to peptidoglycan, rendering the GAC-peptidoglycan linkage resistant to cleavage by mild acid, but sensitive to cleavage by HONO. Importantly, our study reveals that mild acid hydrolysis, following chemical N-acetylation produces nearly quantitative release of GAC with a reducing end sugar (GlcNAc) which is suitable for efficient coupling to free amino groups of target proteins by reductive amination. GAC released by this gentle method provides a much more appropriate substrate for preparation of immunogens because all labile substituents are preserved. Thus, this improved insight into the chemical structure of the GAC-peptidoglycan linkage and the products generated by chemical treatments will guide optimization of conjugation processes with GAC resulting in production of well-defined homogeneous glycoconjugate vaccine.

Previously, similar sensitivity to cleavage by mild acid and HONO was only observed for the WTA of *L. plantarum* AHU 1413 ^20^. We extended the analysis of sensitivity of polysaccharide cleavage by these treatments to other streptococcal species revealing the presence of GlcN residues in the Rha-containing polysaccharide linkers of GBS, *S. mutans* and *S. equi*. In bacteria, enzymatic deacetylation of GlcNAc residues in peptidoglycan, chitin and the exopolysaccharide PNAG is carried out by CE4 family enzymes ^42,47,55,56^. GAS, *S. mutans, S. equi*, and GBS genomes encode two CE4 enzymes, the peptidoglycan deacetylase PgdA and the PplD protein. Using sensitivity to polysaccharide cleavage by mild acid and HONO, we demonstrate that PplD is critical for deacetylation of GlcNAc in the linkage unit of the Rha-containing polysaccharides of GAS, GBS and *S. mutans*. The structural analysis of GAS PplD together with the functional analysis of the site-directed mutants, H105A and D167N, further supports this conclusion. We show that PplD is a metalloenzyme with an active site D-H-H triad coordinating a Zn^2+^ ion. The enzymatic mechanism involves a Zn^2+^-coordinated water molecule acting as a nucleophile, and a conserved histidine residue, H105, acting as a bifunctional acid/base catalyst.

Phylogenetic analysis of PplD homologs in bacterial genomes suggests the occurrence of GlcN 1-phosphate in the peptidoglycan linkage of the polysaccharides in the majority of streptococci except for a few members of the *S. mitis* group that decorate peptidoglycan with choline-containing WTAs. Thus, the presence of PplD in bacterial genomes correlates with the expression of the Rha-containing polysaccharides in these bacteria. Interestingly, the GlcN content of the GAC linkage region varies considerably in the analyzed streptococcal species, from very high in the *S. equi, S. mutans* and GAS strains (about 80-90%) to rather intermediate in GBS A909 (about 30%). Our analysis of the two GAC isoforms with GlcN and GlcNAc linker residues indicates that they are chromatographically and compositionally identical, except for *N*-acetylation of the linkage unit GlcN. Since GAC is presumably synthesized with GlcNAc 1-phosphate at the reducing terminus and it seems unlikely that the LCP transferases catalyzing the transfer of the glycan to cell wall would accept both *N*-acetyl-containing substrates and those devoid of this group, *N*-deacetylation probably occurs after transfer to cell wall. This conclusion is supported by the observation that GlcNAc 1-phosphate-linked polysaccharides, released from GAS saccules by mild acid, appear to be primarily localized to regions of active cell wall synthesis. The simplest explanation is that the nascent GAC chains have not yet had an opportunity to interact with PplD.

At this time, it is not clear what primary advantage deacetylation of linkage unit GlcNAc units might confer on individual streptococci. Introduction of the positive charge into the linkage unit could conceivably stabilize the phosphodiester linkage. However, the presence of the primary amine might also present a vulnerability to hydrolysis by acidified nitrite. In mammals, sodium nitrite serves as a source for generation of nitric oxide (NO), a central biological mediator ^23^. Nitrite is formed via the enzymatic oxidation of NO in the blood and tissues ^57^, and reduction of dietary nitrate by anaerobic bacteria colonizing the oral cavity and gastrointestinal tract ^58^. In the oral cavity, nitrite is actively concentrated by salivary glands reaching up to 2 mM after a dietary nitrate load ^58^. The Streptococcus genus dominates the majority of oral surfaces ^59^. These bacteria produce by-products of fermentation, lactic and acetic acids, that cause a significant decrease in pH level to 5.5 and below. Our unpublished observations indicate that nitrite acidified to pH 5.5 is effective in hydrolysis of rhamnopolysaccharide from *S. mutans*, bacteria associated with dental caries. Since oral microbiota affect many aspects of human health, one may wonder whether streptococci shed the polysaccharides *in vivo*, and whether the products of hydrolysis, free polysaccharide and unmasked peptidoglycan, induce host immune responses and contribute to the complex interplay of oral microbiota.

In *S. iniae*, the PplD homolog was previously identified as a significant virulence factor promoting bacterial infection. The *S. iniae pplD* mutant displayed attenuated virulence in the hybrid striped bass model, decreased survival in whole fish blood and increased sensitivity to lysozyme killing ^43^. We demonstrate that in GAS, PplD and PgdA *N*-deacetylases promote resistance to cationic AMPs (lysozyme, hGIIA, and histones). These AMPs vary from 14 to 21 kDa in size, and attack pathogens by divergent mechanisms ^60^. hGIIA catalyzes the hydrolysis of bacterial phospholipids including phosphatidylglycerol ^61-63^ and is considered a major human host-protective factor against Gram-positive pathogens ^64-66^. Lysozyme catalyzes the hydrolysis of glycosidic bonds of peptidoglycan. The antimicrobial action of histones has been attributed to the disruption of membrane integrity ^67^. Furthermore, similar to histones, lysozyme and hGIIA can insert into negatively charged bacterial membrane forming pores ^52,68-71^. The decreased resistance of the mutants defective in PplD and PgdA to the tested AMPs may be explained by a decrease of the positive charge inside the cell wall. To reach the plasma membrane, AMPs have to traverse the Gram-positive cell envelope. There is substantial evidence that changes in the net charge of cell envelope components, peptidoglycan and teichoic acids, modulate the resistance of bacteria to AMPs ^63,66,72-75^. We previously demonstrated that modification of GAC and the *S. mutans* rhamnopolysaccharide with negatively charged GroP moieties renders GAS and *S. mutans* more sensitive to hGIIA activity ^15^. In this study, we reveal that the GroP modification confers sensitivity of GAS to lysozyme and histone mixture also. Thus, these data support the conclusion that PplD and PgdA contribute to protection of bacteria against AMPs by neutralizing a net negative charge of the cell wall polysaccharides (Fig. 7 **c**).

In sum, the results of this study form a foundation for the design of significantly improved GAS glycoconjugate vaccine based on GAC and provide insight into the function of cell wall modifications in protection of bacteria against cationic AMPs.

## Material and Methods

### Bacterial strains, growth conditions and media

All plasmids, strains and primers used in this study are listed in Supplementary Tables 5 and 6. Streptococcal strains (GAS, GBS, *S. equi* and *S. thermophiles*) were grown in BD Bacto Todd-Hewitt broth supplemented with 1% yeast extract (THY) without aeration at 37°C. *S. mutans* strains were grown on THY agar at 37°C with 5% CO_2_. *E. coli* strains were grown in Lysogeny Broth (LB) medium or on LB agar plates at 37°C. When required, antibiotics were included at the following concentrations: ampicillin and streptomycin at 100 μg mL^-1^ for *E. coli*; erythromycin at 500 μg mL^-1^ for *E. coli* and 5 μg mL^-1^ for streptococci; chloramphenicol at 10 μg mL^-1^ for *E. coli* and 2 μg mL^-1^ for streptococci; spectinomycin at 200 μg mL^-1^ for *E. coli* and 200 μg mL^-1^ for streptococci; kanamycin at 50 μg mL^-1^ for *E. coli* and 300 μg mL^-1^ for streptococci.

### Construction of mutant strains

For construction of the GAS, GBS and *S. mutans* deletion mutants, GAS NZ131, GBS A909 and *S. mutans* Xc chromosomal DNA, respectively, were used as the templates for amplification of DNA fragments encompassing the upstream and downstream regions flanking the gene of interest. The primers used to generate the deletion mutants are listed in Supplementary Table 6. To construct GASΔ*pplD*, the DNA fragments were amplified using two primers pairs JC480/JC481 and JC482/JC483, digested with the restriction enzymes and cloned into pFED760 ^76^. The antibiotic resistance cassette for kanamycin (*aph3A*), amplified from pOskar ^77^ using two primers pairs JC292/JC304, was cloned between the upstream and downstream fragments to generate the plasmid pJC344-kan for selective allelic replacement. The plasmid was electroporated into the GAS NZ131 cells.

To construct GASΔ*pplD*Δ*pgdA*, two DNA fragments were amplified using two primers pairs PplD-BglII-F/PplD-SalI-R and PplD-BamH-F/PplD-XhoI-R (Supplementary Table 2), digested with BglII/SalI and ligated into BglII/SalI-digested pUC19BXspec ^13^. The resultant plasmid was digested with BamHI/XhoI, and used for ligation with the second PCR product that was digested with BamHI/XhoI. The resultant plasmid, pUC19BXspec-*pplD*, was digested with BglII and XhoI to obtain a DNA fragment containing a nonpolar spectinomycin resistance cassette flanked with the *pplD* upstream and downstream regions. This DNA fragment was cloned into BglII/XhoI-digested pHY304 to generate pHY304Δ*pplD*-NZ131. The resultant plasmid was electroporated into the GASΔ*pgdA* ^78^

To construct GASΔ*gacH*, GBSΔ*pplD* and SMUΔ*pplD*, we used a PCR overlapping mutagenesis approach, as previously described ^15^. Briefly, 600-700 bp fragments both upstream and downstream of the gene of interest were amplified with designed primers that contained 16-20 bp extensions complementary to the nonpolar antibiotic resistance cassette (Supplementary Table 6). The nonpolar spectinomycin and chloramphenicol resistance cassettes were PCR-amplified from pLR16T and pDC123, (Supplementary Table 5), respectively, and used for mutagenesis of *pplD* and *gacH*, respectively. Two fragments flanking the gene of interest and the antibiotic resistance cassette were purified using the QIAquick PCR purification kit (Qiagen) and fused by Gibson Assembly (SGA-DNA). A PCR was then performed on the Gibson Assembly sample using primers listed in (Supplementary Table 6) to amplify the fused fragments. The assembled DNA fragment for mutagenesis of *S. mutans pplD* was directly transformed into the *S. mutans* Xc cells by electroporation. The transformants were selected on THY agar containing spectinomycin.

The assembled DNA fragments for mutagenesis of GAS *gacH* and GBS *pplD* were cloned into pHY304 by the use of restriction enzymes to generate the plasmids pHY304Δ*gacH-NZ131* and pHY304Δ*pplD*-GBS, respectively. To plasmids were electroporated into the GAS NZ131 and GBS A909 cells to obtain GASΔ*gacH* and GBSΔ*pplD*, respectively.

A two-step temperature-dependent selection process was used to isolate the desired GAS and GBS mutants ^15^. The transformants were selected on THY agar containing the corresponding antibiotic and screened for sensitivity to erythromycin. Single colonies sensitive to erythromycin were picked. Double-crossover recombination was confirmed by PCR and Sanger sequencing using the primers listed in Supplementary Table 6.

For complementation of GASΔ*pplD*, GASΔ*pplD*Δ*pgdA*, and SMUΔ*pplD*, a promoterless *pplD* gene was amplified from GAS NZ131 using the primers listed in Supplementary Table 6. The PCR product was ligated into pDC123 vector, yielding p*pplD*. To make a catalytically inactive variant of *pplD*, the mutations H105A and D167N were introduced into p*pplD* using Gibson Assembly site-directed mutagenesis (SGI-DNA) with the primers listed in Supplementary Table 6, yielding the p*pplD-H105A*, p*pplD-D167N* and p*pplD-H105A/D167N* plasmids. The plasmids were transformed into GASΔ*pplD*, GASΔ*pplD*Δ*pgdA*, and SMUΔ*pplD*. Chloramphenicol resistant single colonies were picked and checked for presence of p*pplD* by PCR and Sanger sequencing. Complementation of GASΔ*gacH* with WT *gacH* and *gacH-T530A* on pDCerm expression plasmid was conducted as previously described ^15^.

### Construction of the plasmids for expression of ePplD and GFP-AtlA^Efs^

To create a vector for expression of the extracellular domain of PplD (ePplD), the gene was amplified from GAS 5005 chromosomal DNA using the primer pair pplD-NcoI-f and pplD-XhoI-r. The PCR product was digested with NcoI and XhoI, and ligated into NcoI/XhoI-digested pCDF-NT vector. The resultant plasmid, pCDF-PplD, encodes *pplD* fused at the N-terminus with a His-tag followed by a TEV protease recognition site. To construct a vector for expression of GFP-AtlA^Efs^ (pKV1644), 5’ fragment encoding GFP was PCR-amplified from pHR-scFv-GCN4-sfGFP-GB1-NLS-dWPRE using primer pairs sfGFP_BspH and gfp-0799-R, and 3’ fragment encoding a C-terminal cell wall-binding domain of AtlA was amplified from *E. faecalis* V583 DNA using the primer pair gfp-0799-F and 0799-Hind. GFP fusion was constructed using a PCR overlapping mutagenesis approach, digested with BspHI/HindIII and ligated into NcoI/HindIII-digested pRSF-NT vector.

### Crystallization and structure determination of PplD

ePplD was purified for crystallization studies as previously reported for GacH ^15^. Initial screening for crystallization conditions was performed using MCSG I-IV crystallization screens (Anatrace). The optimized crystals were obtained using the vapor diffusion method in 20% PEG3350, 0.2 M zinc acetate, 0.1 M imidazole, pH 8.0. Crystals were transferred to crystallization solution supplemented with 20% glycerol and vitrified in liquid nitrogen prior to data collection. Due to the presence of Zn^2+^ ions in the crystallization solution, the data were collected above Zn anomalous scattering edge at wavelength 1.27 Å at beamline BL9-2 at the Stanford Synchrotron Radiation Laboratory (SSRL). The data were processed and scaled using *XDS* and *XSCALE* ^79^. The structure was solved by single-wavelength anomalous diffraction phasing using an automated *Phaser* SAD pipeline ^80^. The initial zinc positions were found using *SHELXD* ^81^. Following density modification with *Parrot*, the initial model was built using *Buccaneer* ^82,83^. The model was improved using manual rebuilding in *Coot* alternating with refinement by *phenix*.*refine* ^84,85^. The final model had 96% of residues in the favorable region of the Ramachandran plot and 4% in additionally allowed region.

### Lysozyme, hGIIA and histone susceptibility assay

Frozen stocks of GAS variants were used for microbial susceptibility tests with hGIIA, histone mixture from calf thymus (Millipore Sigma, 10223565001) and lysozyme from egg white (Millipore Sigma, L6876). Recombinant hGIIA was produced as described previously ^86^. To make frozen aliquots, GAS strains, cultured overnight in THY medium, were diluted 1:100 into fresh THY medium and allowed to grow to mid-logarithmic phase (OD_600_=0.5). Bacteria were collected by centrifugation, washed twice with PBS and re-suspended in HEPES buffer solution (20 mM HEPES pH 7.5, 2 mM CaCl_2_ and 1% BSA) with 15% glycerol. The aliquots were frozen and stored at −80°C until used in viability assays. Aliquots were thawed and diluted to 1:1000 in HEPES buffer solution. The susceptibility experiments were performed as described previously^75^. Briefly, the AMP (hGIIA, lysozyme and histone mixture) was serially diluted in HEPES solution. Diluted bacterial cultures (10 μl) were mixed with the AMP (10 μl) in a conical 96 well plate. The plate was incubated at 37°C with 5% CO_2_ for 2 hours. After incubation, the bacterial suspensions were diluted with 200 μl PBS and 40 μl was plated on THY agar plates (10 cm) for quantification. After overnight incubation, the CFU were counted. Survival rate was calculated as Survival (% of inoculum) = (counted CFU * 100) / CFU count of original (no AMP added sample). Drop test assay with varying concentrations of lysozyme was conducted as previously described for zinc concentrations ^15^.

### Muramidase activity of lysozyme on GAS

Overnight cultures of the GAS strains were diluted (1:100) into fresh THY broth and grown to an OD_600_ of 0.5. Cell suspension (10 mL) was centrifuged (3,200 g, 10 min) and resuspended with either 1 mg mL^-1^ lysozyme or 62.5 U/ml mutanolysin (Sigma, M9901) or no enzyme (negative control) in sterile PBS (10 mL). After 0 and 18 hours of incubation at 37°C, 0.2 mL was centrifuged (16,000 g, 3 min). The supernatant was transferred to a fresh centrifuge tube and concentrated to 30 μl final volume using SpeedVac vacuum concentrator. Supernatant (5 μl) was spotted on a nitrocellulose membrane. The membrane was blocked for 1 hour with 7% skim milk in PBS/0.1% Tween 20 and incubated with an anti-GAC antibody diluted 1:5000 (Abcam, ab9191). Bound antibodies were detected with a peroxidase-conjugated goat anti-rabbit IG antibody and the Amersham Bioscience ECL (enhanced chemiluminescence) western blotting system. ImageJ software was used to quantify the signal of polysaccharides released.

### Isolation of cell wall and sacculi

Cell wall was isolated from exponential phase cultures by the SDS-boiling procedure as described for *S. pneumoniae* ^87^. Purified cell wall samples were lyophilized and stored at −20°C before the analysis. *S. mutans* and GAS sacculi were obtained by the SDS-boiling procedure followed by 2 washes with 1 M NaCl and 5 washes with water as outlined ^25^. To analyze sensitivity of GAC cleavage to chemical treatments, the GAS sacculi were additionally treated with 5 mg mL^-1^ trypsin (Sigma, T-8253) at 37°C overnight, sedimented (32,000 g for 15 min), and washed 4 times by resuspension in water. This treatment removes binding of the GAS sacculi to the microcentrifuge plastic tubes. Sacculi were resuspended in water and stored at 4 C before the analysis.

### HONO deamination

Polysaccharide was released from sacculi or highly enriched cell wall preparations by HONO deamination essentially as described ^22^ with minor modification. Briefly, reactions contained sacculi (OD_600_ = 3.4) or isolated cell walls (2-5 mg mL^-1^), in 0.2 N Na Acetate pH 4.5 and 1.5 M NaNO_2_ (added in three equal portions over a period of 90 minutes) at room temperature.

Following deamination, the reaction was stopped by the addition of one equivalent of ethanolamine and centrifuged (60,000 g, 10 min). The supernatant was removed and transferred to an Amicon Ultra-0.5 mL (3,000 MWCO) spin filter. The pellet was washed two times by resuspension in water (1/2 of the original reaction volume), re-sedimented, and the washings were combined with the initial supernatant. The washed pellet was re-suspended in water and reserved for analysis, or discarded. The released material was desalted by several sequential cycles of dilution with water and re-concentration on the spin filter, recovered in a convenient volume of water and either reserved for analysis or purified further by chromatography on Bio Gel P150 and ion exchange on DEAE-Toyopearl.

### Mild acid hydrolysis

Mild acid hydrolysis was performed, either with or without prior *N*-acetylation, as previously described for WTA of *L. plantarum* ^20^ with some modifications. Chemical N-acetylation reactions contained sacculi (OD_600_= 3.4) or isolated cell walls (2-5 mg/ml), resuspended in 1 mL of saturated NaHCO_3_, and 2 % acetic anhydride. After 1 h on ice, the reactions were allowed to warm to room temperature overnight. After incubation at room temperature overnight, the reactions were diluted with 2 volumes of water and sedimented (50,000 x g, 10 min). The insoluble residue was washed three times by resuspension with 3 mL water followed by sedimentation (50,000 x g, 10 min). For mild acid hydrolysis, sacculi or cell wall material was re-suspended in 0.02 N HCl and heated to 100°C for 20 min. The reactions were cooled on ice, neutralized by the addition of 1 N NaOH (1 equivalent) and sedimented (50,000 x g, 10 min). The supernatant fraction was removed and reserved, and the pellet was re-suspended in water and re-sedimented. The supernatant fractions were combined and either analyzed or purified further. The pellet fractions were resuspended in an equal volume of water and reserved for further analysis for rhamnose content or used for fluorescent microscopy.

### Fluorescent and differential interference contrast (DIC) microscopy

GAS sacculi obtained by the SDS-boiling procedure were subjected to mild acid hydrolysis or deamination with HONO as outlined above. Untreated or treated sacculi were resuspended in PBS to an OD_600_=0.3, and incubated with 19.8 μg mL^-1^ GFP-AtlA^Efs^ for 30 min at room temperature. The samples were washed three times with water, dried at room temperature, and mounted on a microscope slide with ProLong Glass Antifade (Invitrogen). Samples were imaged on a Leica SP8 equipped with 100X, 1.44 N.A. objective and DIC optics.

### Chromatography of GAC

GAC, released by either deamination with HONO or by mild acid treatment, was partially purified by size-exclusion chromatography (SEC) on a 1 × 18 cm column Bio Rad BioGel P150 equilibrated in 0.2 N Na acetate, pH 3.7, 0.15 M NaCl. Following SEC, GAC was further fractionated by ion-exchange chromatography on a 1 × 18 cm column of DEAE-Toyopearl (TosoHaas) equilibrated in 10 mM HEPES, pH 8. After elution with two column volumes, DEAE-bound glycans were eluted with an 80 mL gradient (0-0.5 M) of NaCl.

### Phosphate assays

Total phosphate content of SCCs was determined by the malachite green method following digestion with perchloric acid, as previously described ^15^.

### Modified anthrone assay

Total Rha content was estimated using a minor modification of the anthrone procedure as previously described ^15^. Rha concentrations were estimated using L-Rha standard curves.

### Analysis of glycosyl composition by gas chromatography-mass spectrometry (GC-MS)

Saccharide compositions were determined as either trimethylsilyl (TMS) derivatives of O-methyl glycosides, following methanolysis or as alditol acetates following hydrolysis with 2 N TFA and peracetylation, as described below ^88^. Methanolic HCl (1 N) was prepared by the careful dilution of the required amount of acetyl chloride into rapidly stirred, anhydrous methanol at 0°C. TMS-methyl glycoside samples contained 10 nmole inositol as internal standard and were dried thoroughly from absolute ethanol (3 times), under dry nitrogen gas in glass 13 × 100 screw cap tubes with teflon-lined caps. Dried samples were dissolved in 0.2 mL 1 N methanolic HCl and heated to 100°C, 3 h. Following methanolysis, samples were neutralized with 5-10 mg AgCO_3_ and re-*N*-acetylated with 0.1 mL acetic anhydride overnight. Re-*N*-acetylated methyl glycosides were recovered from the solid residue in 1 mL of methanol, dried thoroughly with toluene 2 times, to azeotrope the acetic anhydride, and trimethysilylated with 25 μL TMS-HT (TCI America) at 80°C, 30 min. TMS-methyl glycosides were dried briefly with addition of toluene under dry nitrogen gas, redissolved in 0.1 mL hexane and analyzed by gas chromatography/chemical ionization (methane) mass spectrometry on a Thermo Scientific Trace 1310 gas chromatograph, 0.25 mm x 15 m DB-1701 (J&W Scientific) capillary column, and an ISQ LT single quadrupole mass spectrometer in selective ionization mode (SIM) monitoring appropriate specific high molecular ions. Ion source temperature was 200°C and transfer line temperature was 240°C. Samples (1 μL) were injected in split mode (split flow 8 mL/min, split ratio 16:1) and chromatographed with He carrier gas (0.5 mL/min) with a temperature program of 100°C for 1 min, and 10°C/min to 280°C with a final time of 2 min and detected in positive ion mode. The unknown TMS-sugars co-chromatographed with derivatives of authentic sugar standards and their respective full scan mass spectra contained [MH]^+^, [MH-16]^+^, [MH-32]^+^, [MH-90]^+^, [MH-122]^+^ and [MH-180]^+^ as the only high molecular weight ions. Specific ions for selective ion monitoring were: TMS_3_-*O*-methyl-rhamnose (M/Z = 363 [MH-32]^+^), TMS_4_-*O*-methyl-glucose (M/Z = 361 [MH-122]^+^), TMS_6_-inositol (M/Z = 433 [MH-180]^+^) and TMS_3_-*O*-methyl-GlcNAc (M/Z = 420 [MH-32]^+^).

Sodium borohydride-reduced, intact mild acid-or HONO-released polysaccharides, were fractionated by SEC on BioGel P150 and ion-exchange chromatography on DEAE-Toyopearl, and analyzed for reducing end sugars as alditol acetates. Polysaccharides were concentrated and desalted over Amicon Ultra-0.5, 3,000 Da cut-off, centrifugal filters, supplemented with 1 nmol mannitol internal standard and hydrolyzed in 2 N TFA, 120°C, 1 hr. Samples were dried under air with addition of 1-propanol 3 times and peracetylated with 0.2 mL pyridine/acetic anhydride (1:1), 100°C, 30 min. Samples were diluted with 2 mL CHCl_3_, partitioned with 2 mL water 3 times and dried under a stream of dry nitrogen gas with addition of toluene at room temperature. Alditol acetate analysis was carried out as described above for TMS-methyl glycosides except that the temperature program was 120°C, 1 min, and 5°C per minute to 250°C, with a final time of 2.5 min. Full scan spectra of the alditol acetates contained [MH-60]^+^ as the only high molecular weight ion detected. Chromatograms were prepared by selected ion monitoring using: Ac_4_-2,5-anhydromannitol (M/Z = 273 [MH-60]^+^), Ac_6_-mannitol (M/Z = 375 [MH-60]^+^) and Ac_6_-GlcNAcitol (M/Z = 374 [MH-60]^+^).

Quantities of component sugars were calculated from GC areas after normalization to the internal standard and response factor corrections. Response factors were calculated from standard mixtures of sugars and internal standard using the formula: Rf = (A_x_/A_IS_)/(C_x_/C_is_) in which, A_x_ is GC area of analyte, A_IS_ is GC area of internal standard, C_x_ is amount of analyte and C_is_ is amount of internal standard. Amounts of sugars were calculated using the formula: C_x_ = C_is_×(A_x_/A_IS_)/Rf. In this analysis, an exponential increase in response factor associated with increasing amounts of analyte relative to internal standard was observed. To correct for this change in response factor, standard curves of increasing amounts of analytes to a constant quantity of internal standard were prepared and used to estimate the appropriate response factor.

### NMR spectroscopy

Cell wall isolated from GAS 5005 was used for purification of GAC for NMR analysis ^15^. NMR experiments on the GAC polysaccharide, ∼7 mg in D_2_O (99.96 %) pD ∼8, were acquired at 323.2 K unless otherwise stated using the following Bruker spectrometers: AVANCE III 700 MHz equipped with a 5 mm TCI Z-Gradient Cryoprobe (^1^H/^13^C/^15^N), AVANCE III 600 MHz equipped with a 5 mm TXI inverse Z-Gradient probe (^1^H/^31^P/^13^C), Avance III HD 800 MHz equipped with a 3 mm TCI cryoprobe and an Avance III HD 600 MHz equipped with a 5 mm BBI probe. Chemical shifts are reported in ppm using internal sodium 3-trimethylsilyl-(2,2,3,3-^2^H_4_)-propanoate (TSP, δ_H_ 0.00 ppm), external 1,4-dioxane in D_2_O (δ_C_ 67.40 ppm) and 2% H_3_PO_4_ in D_2_O (δ_P_ 0.00 ppm) as references.

^1^H,^1^H-TOCSY NMR experiments were recorded with mixing times of 10, 30, 60, 90 and 120 ms. ^1^H,^1^H-NOESY experiments ^89^ were collected with mixing times of 100 and 200 ms at 700 MHz and at 800 MHz. Multiplicity-edited ^1^H,^13^C-HSQC experiments ^90^ were carried out with an echo/antiecho-TPPI gradient selection with and without decoupling during the acquisition. ^1^H,^13^C-HSQC-TOCSY experiments were acquired using the MLEV17 sequence for homonuclear Hartman-Hahn mixing with durations of 20, 40, 80 and 120 ms, an echo/antiecho-TPPI gradient selection and heteronuclear decoupling during acquisition. ^1^H,^13^C-HMBC experiment and band-selective ^1^H,^13^C-CT-HMBC were run with a nominal long-range *J* value of 8 Hz. ^1^H,^31^P-Hetero-TOCSY experiments ^27^ were collected using a DIPSI2 sequence with mixing times of 50 and 80 ms. ^1^H,^31^P-HMBC experiments were recorded using an echo/antiecho gradient selection using 25, 50 and 90 ms delays for *J* evolution, 8k and 512 points in the *F*_2_ and *F*_1_ dimensions, respectively, a nominal value for long-range ^n^*J*_CH_ of 8 Hz and a three-fold low-pass J-filter to suppress one-bond correlations. The 3D ^1^H,^13^C,^31^P ^91^ spectra were obtained using echo/anti-echo gradient selection and constant time in *t*_2_ with a nominal value of ^n^*J*_CP_ of 5 Hz and without multiplicity selection.

Non-uniform sampling (NUS) was employed for sparse sampling of 2D NMR experiments with a 50% or 25% coverage, thereby facilitating an increase in the number of increments in the indirect dimension by a factor of 2 or 4, respectively, with overall equal measurement time. The Multi-Dimensional Decomposition (MDD-NMR) ^92-94^ algorithm was employed to reconstruct the FIDs. The spectra were processed and analyzed using TopSpin 4.0.1 software (Bruker BioSpin).

### Statistical analysis

Unless otherwise indicated, statistical analysis was carried out on pooled data from at least three independent biological repeats. Quantitative data was analyzed using either one-way or two-way ANOVA with either Tukey’s or Bonferroni multiple comparison test. Symbols and error bars represent the mean and S.D., respectively. A *p*-value equal to or less that 0.05 was considered statistically significant.

## Data availability

Atomic coordinates and structure factors of PplD crystal structure have been deposited to the Protein Data Bank with accession code 6DQ3. All data generated during this study are included in the article and Supplementary information files or will be available from the corresponding author upon reasonable request.

## Acknowledgments

The authors thank Dr. John F. Timoney (University of Kentucky) for providing *S. equi*, Dr. Jeffrey M. Bosken and Dr. Edward D. Hall (University of Kentucky) for the use of the Thermo Fisher Scientific GC-Mass Spectrometer, Dr. Catalina Velez-Ortega (University of Kentucky) for the access to Leica SP8 confocal microscope, and Dr. Peter H. Spielmann (University of Kentucky) for helpful discussions. This work was supported by NIH grants R01 AI143690 from the NIAID and R01 DE028916 from the NIDCR and (to NK), the Swedish Research Council (no. 2017-03703) and The Knut and Alice Wallenberg Foundation (GW). The Swedish NMR Centre at University of Gothenburg is acknowledged for support. Use of the Stanford Synchrotron Radiation Lightsource, SLAC National Accelerator Laboratory, is supported by the U.S. Department of Energy, Office of Science, Office of Basic Energy Sciences under Contract No. DE-AC02-76SF00515. The SSRL Structural Molecular Biology Program is supported by the DOE Office of Biological and Environmental Research, and by the National Institutes of Health, National Institute of General Medical Sciences (P30GM133894). The funders had no role in study design, data collection and interpretation, or the decision to submit the work for publication.

## Author contributions

JSR, AR, GL, MJF, KVK, GW, and NK designed the experiments. JSR, PP, AAP, CWK, AG, MMR, GL, GW and NK performed functional and biochemical experiments. JL and KVK carried out X-ray crystallography and structure analysis. AR and GW performed NMR studies. JCC, KVK and NK constructed plasmids and isolated mutants. SZ performed microscopy analysis. JSR, AR, KVK, GW, and NK analyzed the data. NK wrote the manuscript with contributions from all authors. All authors reviewed the results and approved the final version of the manuscript.

