## Supplementary Information for "PplD is a de-N-acetylase of the cell wall linkage unit of streptococcal rhamnopolysaccharides"

**Supplementary Table 1**. $^{1}H$, $^{13}C$ and $^{31}P$ NMR chemical shifts (ppm) at 50 °C of the linker region from Group A Carbohydrate and inter-residue correlations from $^{1}H$,$^{1}H$-NOESY, $^{1}H$,$^{13}C$-HMBC and $^{1}H$,$^{31}P$-HMBC experiments.

| **Sugar residue** | **$^{1}H/^{13}C$** | | | | | | | | **$^{31}P$** | **Inter-residue correlation** | |
|---|---|---|---|---|---|---|---|---|---|---|---|
| | 1 | 2 | 3 | 4 | 5 | 6 | α | β | $PO_4$ | NOE | HMBC[b] |
| →3)-β-L-Rha*p*-(1→ | 4.89 {164}[a] | 4.18 | 3.65 | 3.50 | 3.43 | 1.33 | | | | H4, **GlcN** | C4, **GlcN** |
| | 101.6 | 71.4 | 81.4 | 72.1 | 72.9 | 17.6 | | | | | |
| →4)-α-D-Glc*p*N(1-*P* | 5.54 {177} | 2.89 | 3.94 | 3.69 | 3.77 | ~3.88 | | | -1.35 | | H1, **GlcN**<br>H2, **GlcN** |
| | 96.4 | 56.0 | 72.9 | 77.7 | 73.6 | 61.5 | | | | | |
| *P*-6)-β-D-Mur*p*NAc-(1→ | 4.57 {167} | 3.85 | 3.64 | 3.93 | 3.47 | 4.10, 4.20 | 4.37 | 1.41 | | | H6a, **MurNAc**<br>H6b, **MurNAc** |
| | 102.4 | 55.9 | 80.4 | 75.9 | 75.0 | 64.4 | 79.2 | 19.0 | | | |

[a] $^{1}J_{CH}$ values are given in Hertz in braces; [b] Refers to $^{1}H$,$^{13}C$-HMBC or $^{1}H$,$^{31}P$-HMBC experiments.

**Supplementary Table 2.** Binding of GFP-AtlA$^{Efs}$ to the GAS sacculi.

| | Total number of cells counted | Number of cells with GFP-AtlA$^{Efs}$ bound to the poles | Number of cells with GFP-AtlA$^{Efs}$ bound to the whole cell surface | Number of dividing cells | Number of dividing cells with GFP-AtlA$^{Efs}$ bound to the septal regions |
|---|---|---|---|---|---|
| Untreated | 127 | 34 (26%)[a] | 0 | 41 | 8 (19%)[c] |
| Treated with mild acid | 183 | 129 (70%) | 0 | 70 | 65 (92.9%) |
| Treated with nitrous acid | 117 | 117 (100%) | 117 (100%)[b] | 47 | 117 (100%) |

[a] The percentage of cells with GFP-AtlA$^{Efs}$ attached to the poles.

[b] The percentage of cells with GFP-AtlA$^{Efs}$ attached to the whole cell surface.

[c] The percentage of dividing cells with GFP-AtlA$^{Efs}$ attached to the septal regions.

**Supplementary Table 3.** Data collection and refinement statistics for PplD structure.

| | PDB: 6DQ3[1] |
|---|---|
| **Data collection** | |
| Space group | $P2_12_12_1$ |
| Cell dimensions | |
| *a*, *b*, *c* (Å) | 42.16, 78.43, 138.07 |
| α, β, γ (°) | 90, 90, 90 |
| | *Peak* |
| Wavelength (Å) | 1.2700 |
| Resolution (Å) | 39.69–1.78 (1.83–1.78)[2] |
| $R_{sym}$ or $R_{merge}$ | 0.065 (1.126)[3] |
| $I / \sigma I$ | 13.04 (1.09) |
| Completeness (%) | 99.3 (97.5) |
| Redundancy | 3.8 (3.3) |
| **Refinement** | |
| Resolution (Å) | 39.69–1.78 |
| No. reflections (total / free) | 84025 / 4297 |
| $R_{work}$ / $R_{free}$ | 0.180 / 0.216 |
| No. atoms | |
| Protein | 3597 |
| Ligand/ion | 37 |
| Water | 296 |
| *B*-factors | |
| Protein | 35.1 |
| Ligand/ion | 39.6 |
| Water | 38.1 |
| Wilson *B* | 33.5 |
| R.m.s deviations | |
| Bond lengths (Å) | 0.004 |
| Bond angles (°) | 0.685 |

[1]One crystal was used for data collection.
[2]Values in parentheses are for highest-resolution shell.
[3]Friedel pairs are treated as separate reflections.

**Supplementary Table 4.** Structural homologs of GAS PplD.

| PDB ID | Z score | r.m.s.d. | Number of aligned residues | Total number of residues | Sequence identity, % | Protein function | Reference |
|---|---|---|---|---|---|---|---|
| 4hd5 | 30.3 | 1.4 | 209 | 316 | 31 | *Bacillus cereus*, polysaccharide deacetylase | [1] |
| 4v33 | 30.1 | 1.5 | 211 | 316 | 30 | *Bacillus anthracis*, polysaccharide deacetylase-like protein | [2] |
| 6go1 | 29.2 | 1.7 | 211 | 318 | 27 | *Bacillus anthracis*, polysaccharide deacetylase-like protein | [3] |
| 4wcj | 26.7 | 2.1 | 208 | 233 | 24 | *Ammonifex degensii*, polysaccharide deacetylase | [4] |
| 4u10 | 26.3 | 2.7 | 221 | 264 | 19 | *Aggregatibacter actinomycetemcomitans*, poly-β-1,6-N-acetyl-D-glucosamine N-deacetylase | NP[a] |
| 3vus | 25.8 | 2.3 | 215 | 263 | 21 | *Escherichia coli*, poly-β-1,6-N-acetyl-D-glucosamine N-deacetylase | [5] |
| 4f9d | 25.7 | 2.5 | 219 | 592 | 21 | *Escherichia coli*, poly-β-1,6-N-acetyl-D-glucosamine N-deacetylase | [6] |
| 5bu6 | 22.6 | 2.5 | 197 | 264 | 19 | *Bordetella bronchiseptica*, poly-β-1,6-N-acetyl-D-glucosamine N-deacetylase | [7] |
| 2c1g | 11.7 | 2.4 | 140 | 384 | 26 | *Streptococcus pneumoniae*, peptidoglycan GlcNAc deacetylase | [8] |

[a]NP – no publication.

**Supplementary Table 5.** Bacterial strains and plasmids.

| Strain or plasmid | Description [a] | Reference |
|---|---|---|
| ***Bacteria*** | | |
| GAS NZ131 | M49-serotype strain | [9] |
| GAS 5005 | M1T1-serotype strain | [10] |
| *S. mutans Xc* | Serotype *c* strain | [11] |
| *S. equi* | *S. equi* subsp. *equi* CF32 was isolated from an equine submandibular abscess. | [12] |
| *S. agalactiae* A909 | Human clinical isolate. Serotype Ia strain, ST-7 | ATCC |
| *S. agalactiae* COH1 | Clinical isolate obtained from an infected newborn with sepsis. Serotype III strain, ST-17 | [13] |
| *S. thermophilus* LMG 18311 | Isolated from yogurt manufactured in the United Kingdom | ATCC |
| GASΔ*pgdA* | *pgdA* deletion mutant in the GAS NZ131 strain background (has a nonpolar kanamycin resistance cassette replacing *pgdA*), $Kan^R$ | [14] |
| GASΔ*pplD* | *pplD* deletion mutant in the GAS NZ131 strain background (has a nonpolar kanamycin resistance cassette replacing *pplD*), $Kan^R$ | This study |
| GASΔ*pplD*Δ*pgdA* | *pplD* deletion mutant in the GASΔ*pgdA* strain background (has nonpolar spectinomycin and kanamycin resistance cassettes inserted in *pplD* and *pgdA*, respectively), $Spec^R$, $Kan^R$ | This study |
| GASΔ*pplD*:p*pplD* | GASΔ*pplD* is complemented with p*pplD* carrying GAS WT *pplD,* $Kan^R$, $Cam^R$ | This study |
| GASΔ*pplD*:p*pplD*-*H105A* | GASΔ*pplD* is complemented with p*pplD* carrying a catalytically inactive variant of GAS *pplD*-H105A*,* $Kan^R$, $Cam^R$ | This study |
| GASΔpplD:p*pplD*-D167N | GASΔ*pplD* is complemented with p*pplD* carrying a catalytically inactive variant of GAS *pplD*-D167N, $Kan^R$, $Cam^R$ | This study |
| GASΔ*pplD*:p*pplD*--*H105A/D167N* | GASΔ*pplD* is complemented with p*pplD* carrying a catalytically inactive variant of GAS *pplD*-H105A/D167N*,* $Kan^R$, $Cam^R$ | This study |
| GASΔ*pplD*Δ*pgdA*:p*pplD* | GASΔ*pplD*Δ*pgdA* is complemented with p*pplD* carrying GAS WT *pplD*, $Spec^R$, $Kan^R$, $Cam^R$ | This study |
| GASΔ*gacH* | *gacH* deletion mutant in the GAS NZ131 strain background (has a nonpolar chloramphenicol resistance cassette inserted in *gacH*), $Cam^R$ | This study |
| GASΔ*gacH*:p*gacH* | GASΔ*gacH* is complemented with p*gacH*_erm carrying WT *gacH*, $Cam^R$, $Erm^R$ | This study |
| GASΔ*gacH*:p*gacH*-T530A | GASΔ*gacH* is complemented with p*gacH*-T530A *c*arrying a catalytically inactive variant of *gacH*, $Cam^R$, $Erm^R$ | This study |
| GBSΔ*pplD* | *pplD* deletion mutant in the GBS A909 strain background (has a nonpolar spectinomycin resistance cassette replacing *pplD*), $Spec^R$ | This study |
| GBSΔ*pplD*:p*pplD* | GBSΔ*pplD* is complemented with p*pplD* carrying GAS WT *pplD,* $Spec^R$, $Cam^R$ | This study |

| | | |
|---|---|---|
| SMUΔ*pplD* | *pplD* deletion mutant in the *S. mutans* Xc strain background (has a nonpolar spectinomycin resistance cassette replacing *pplD*), Spec$^R$ | This study |
| SMUΔ*pplD:*p*pplD* | SMUΔ*pplD* is complemented with p*pplD* carrying GAS WT *pplD,* Spec$^R$, Cam$^R$ | This study |
| SMUΔ*pplD:*p*pplD*-H105A | SMUΔ*pplD* is complemented with p*pplD* carrying a catalytically inactive variant of GAS *pplD*-H105A*,* Spec$^R$, Cam$^R$ | This study |
| SMUΔ*pplD:*p*pplD*-D167N | SMUΔ*pplD* is complemented with p*pplD* carrying a catalytically inactive variant of GAS *pplD*-D167N, Spec$^R$, Cam$^R$ | This study |
| SMUΔ*pplD:*p*pplD*-H105A/D167N | SMUΔ*pplD* is complemented with p*pplD* carrying a catalytically inactive variant of GAS *pplD*-H105A/D167N*,* Spec$^R$, Cam$^R$ | This study |
| ***Escherichia coli*** | | |
| DH5α | *E. coli* cells used for cloning | Invitrogen |
| Rosetta (DE3) | *E. coli* cells used for protein expression; Cam$^R$ | Novagen |
| ***Plasmids*** | | |
| pHY304 | A temperature sensitive *E.coli-Streptococcus* shuttle vector, Erm$^R$ | [15] |
| pHY304Δ*gacH*-NZ131 | Derivative of pHY304 expressing a nonpolar chloramphenicol resistance cassette flanked with GAS NZ131 *gacH* 5' and 3' regions, Cam$^R$, Erm$^R$ | This study |
| pUC19BXspec | Derivative of pUC19BX expressing *aadA* (a nonpolar spectinomycin) with own rbs, Amp$^R$ | [16] |
| pUC19BXspec-*pplD* | Derivative of pUC19BXspec expressing *aadA* flanked with *pplD* 5' and 3' regions, Amp$^R$ | This study |
| pHY304Δ*pplD*-NZ131 | Derivative of pHY304 expressing a nonpolar spectinomycin resistance cassette flanked with GAS NZ131 *pplD* 5' and 3' regions, Spec$^R$, Erm$^R$ | This study |
| pHY304Δ*pplD*-GBS | Derivative of pHY304 expressing a nonpolar spectinomycin resistance cassette flanked with GBS A909 *pplD* 5' and 3' regions, Spec$^R$, Erm$^R$ | This study |
| pFED760 | A temperature sensitive *E.coli-Streptococcus* shuttle vector, Erm$^R$ | [17] |
| pOskar | Vector encoding kanamycin resistance cassette, Kan$^R$ | [18] |
| pLR16T | Vector encoding spectinomycin resistance cassette, Spec$^R$ | [19] |
| pDC123 | *E. coli-streptococcus* shuttle vector, JS-3 replicon, chloramphenicol resistance cassette. Cam$^R$ | [20] |
| p*pplD* | pDC123 derived plasmid expressing *pplD.* Cam$^R$ | This study |
| p*pplD*-H105A | pDC123 derived plasmid expressing *pplD*-H105A, Cam$^R$ | This study |
| p*pplD*-D167N | pDC123 derived plasmid expressing *pplD*-D167N, Cam$^R$ | This study |
| p*pplD*-H105A/D167N | pDC123 derived plasmid expressing *pplD*-H105A/D167N, Cam$^R$ | This study |
| pDCerm | pDC123 derivative with Erm (Erm of Tn916ΔE) replacing chloramphenicol resistance cassette, Erm$^R$ | [21] |
| p*gacH*_erm | pDCerm derived plasmid expressing *gacH,* Erm$^R$ | [22] |

| | | |
|---|---|---|
| p*gacH*-T530A | p*gacH*_erm derived plasmid expressing a catalytically inactive of variant of *gacH,* $Erm^R$ | [22] |
| pRSF-NT | A modified pRSF-Duet1 (Novagen) vector that allows the creation of N-terminus His-tagged proteins with a TEV protease cleavage site, $Kan^R$ | [23] |
| pKV1644 | pRSF-NT derived plasmid for expression of GFP-$AtlA^{Efs}$. A C-terminal cell wall-binding domain of AtlA from *E. faecalis* fused with GFP at the N-terminus. GFP has a His-tag followed by a TEV protease recognition site at the N-terminus, $Kan^R$ | This study |
| pCDF-NT | A modified pCDF-Duet1 (Novagen) vector for expression of the N-terminus His-tagged proteins with a TEV protease cleavage site, $Str^R$ | [24] |
| pCDF-PplD | pCDF-NT derived plasmid expressing ePplD fused with N-terminal TEV protease cleavable His-tag, $Str^R$ | This study |

[a] Antibiotic resistance markers: $Erm^R$, erythromycin; $Kan^R$, kanamycin; $Spec^R$, spectinomycin; $Cam^R$, chloramphenicol, $Amp^R$, ampicillin, $Str^R$, streptomycin

**Supplementary Table 6.** Primers

| Primer | Sequence[a,b] | Genetic manipulations |
|---|---|---|
| JC480 | CATGGAATTCCCAGATTGAAGCACCAACC | GAS *pplD* deletion with a nonpolar kanamycin resistance cassette |
| JC481 | CATGACGCGTTACTTGCTCCTTTTTTTGATAT | |
| JC482 | CATGACGCGTCATCAATTTTTTTATCATTCCAAA | |
| JC483 | CATGGAATTCAACTTATTCAGTTCAAGACCTG | |
| JC292 | GCATGACGCGTATGGCTAAAATGAGAATATCACC | |
| JC304 | GCATGACGCGTCTAAAACAATTCATCCAGTAAAATATAA | |
| PplD-BglII-F | GCGTAAGATCTGTGATCATGAATCCATTCTAG | GAS *pplD* deletion with a nonpolar spectinomycin resistance cassette |
| PplD-SalI-R | CGCTGCGTCGACGGGTTGACAATCAAATTAGC | |
| PplD-BamH-F | CGTCTGGATCCCAGTATGATTGATTTCTACAAC | |
| PplD-XhoI-R | GCGCGCTCGAGGACGTCTCCAAGAATAACAGC | |
| GacHNZ-BamH-F2 | CGTCTGGATCCCGTTGGTCCCGCTAAAGCAATG | GAS NZ131 *gacH* deletion with a nonpolar chloramphenicol resistance cassette |
| cm-gacHNZ-R1 | **GTGAATTTAGGAGGC**CGTATATGATTGTTGCAAATATG | |
| gacHNZ-cam-F1 | CAACAATCATATACG**GCCTCCTAAATTCACTTTAG** | |
| Cm-gacHNZ-F2 | **AATATGAGATAATGC**GGTGATGAAGCATTGCTAGG | |
| gacHNZ-Cm-R2 | CAATGCTTCATCACC**GCATTATCTCATATTATAAAAG** | |
| GacHNZ-XhoI-R | GCGCGCTCGAGCATAAGTCCCGCAGTTGTGC | |
| PplD-Smu-F | CCGCTTCTTGTGCTATAATAAG | *S. mutans pplD* deletion with a nonpolar spectinomycin resistance cassette |
| Spec-PplD-Smu-r1 | **CTCACTATTTTGGTCGAC**CTTTAGCTCTGCAGCCTGCG | |
| PplD-Smu-Spec-f1 | CAGGCTGCAGAGCTAAAG**GTCGACCAAAATAGTGAGGAG** | |
| Spec-PplD-Smu-f2 | **AAAAATTATAAGGATCC**CAGCAATAGCTTATCCAGCGG | |
| PplD-Smu-Spec-r2 | CTGGATAAGCTATTGCTG**GGATCCTTATAATTTTTTAATCTG** | |
| PplD-Smu-XhoI-R | GGAAGAAATATGTTTGGAAGC | |
| PplD-GBS-BamHI-F | CGTCTGGATCCGTATTTATCTCTGTCACAAAG | GBS *pplD* deletion with a nonpolar spectinomycin resistance cassette |
| Spec-PplD-GBS-R1 | **CTCACTATTTTGGTCGAC**GCTAAGAAGTAATATCCTTCC | |
| PplD-GBS-spec-F1 | GATATTACTTCTTAGC**GTCGACCAAAATAGTGAGGAG** | |
| Spec-PplD-GBS-F2 | **AAAATTATAAGGATC**GATGATGGAAATGCCGATTTC | |
| PplD-GBS-spec-R2 | GGCATTTCCATCATC**GATCCTTATAATTTTTTAATCTG** | |
| PplD-GBS- | GCGCGCTCGAGTTGGCAGTGATTGCTATTG | |

| | | |
|---|---|---|
| XhoI-F | | |
| sfGFP_BspH | GAGATCATGAGCAAAGGAGAAGAACTTTTCAC | Construction of pKV1644 |
| gfp-0799-R | **GCCCCCAGTATT**ACCACCTCCACCTTTGTAGAGC | |
| gfp-0799-F | ACAAAGGTGGAGGTGGT**AATACTGGGGGCGGAAC** | |
| 0799-Hind | CAGAAGCTTAACCAACTTTTAAAGTTTGACCAATATAAATTG | |
| PplD-NcoI-f | CGTGAGCCATGGAAACACCTGTCAAGATCCC | Construction of pCDF-PplD |
| PplD-XhoI-r2 | GCGCGCTCGAGTTATGGTTCCATTGTTTGTAAAAG | |
| PplD-HindIII-f | CGCGCAAGCTTGTTACAATAGAGGTACTTATATC | Construction of p*pplD* |
| PplD-BamHI-r2 | CGTCTGGATCCTTATGGTTCCATTGTTTGTAAAAG | |
| PplD-GAS-check-f | CTTGGACCAAGCAAGAAACAC | Verification of GASΔ*pplD* |
| PplD-GAS-check-r | CTTCCCATTTCGGTTAGTCC | |
| gacHNZ-check-F | CTGTGGATAGTTTTACTTGTC | Verification of GASΔ*gacH* |
| gacHNZ-check-R | CAACAGAAATAATTGTTCCC | |
| PplD-SMU-check-f | CCCATCCTGATTTATCTGCTTC | Verification of SMUΔ*pplD* |
| PplD-SMU-check-r | CCATCGTTAGCACTAGCTAGGC | |
| pplDGBS-check-f | CCATGCTGTTCATGTTATGG | Verification of GBSΔ*pplD* |
| pplDGBS-check-r | GCATTTGTTGAACGTTGAGG | |
| H105A_F | CCCAATTTTAATGTATGCTGCTATTCATGTAATGTCC | H105A mutation in PplD, construction of p*pplD*-H105A |
| H105A_R | GGACATTACATGAATAGCAGCATACATTAAAATTGGG | |
| D167N_F | CGCCCTTAGACACGCTACCGCTACCAGATGGTAGAGCAAC | D167N mutation in PplD, construction of p*pplD*-D167N and p*pplD*-H105A/D167N |
| D167N_R | CAATCATACTGTCATTAAATGTTAGCCATACAAC | |

[a] Restriction sites are underlined.
[b] Extensions complementary to the antibiotic resistance cassettes or GFP are in bold.

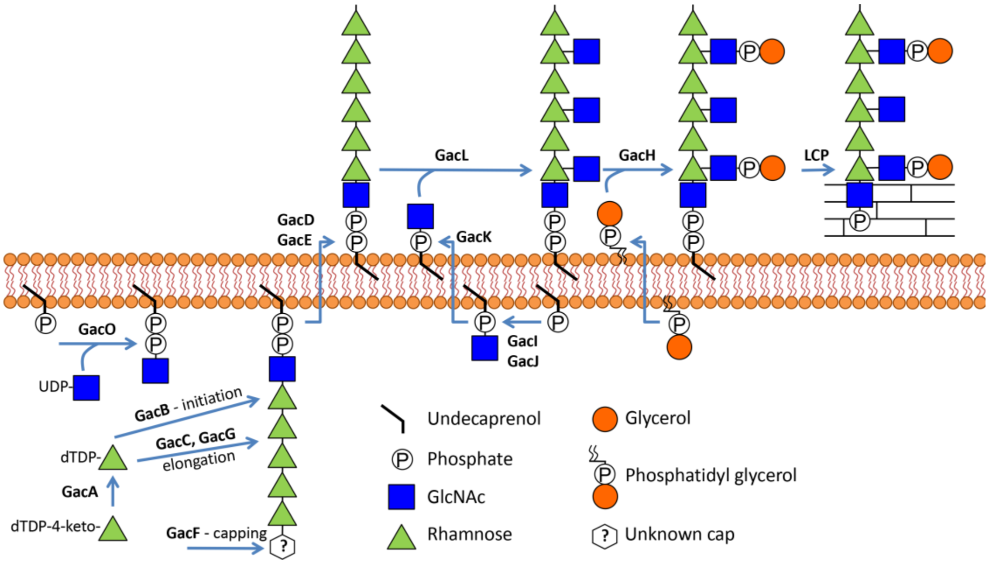

**Supplementary Fig. 1. A proposed mechanism of GAC biosynthesis.**

GAC biosynthesis is initiated on the inner leaflet of the plasma membrane by GacO which transfers GlcNAc from UDP-GlcNAc to undecaprenyl phosphate (Und-P) producing GlcNAc-P-P-Und [16]. The lipid serves as a membrane-anchored acceptor for GacB-mediated transfer of the first sugar residue, L-Rha, from TDP-β-L-Rha to form Rha-GlcNAc-P-P-Und [25]. The GacC, GacF and GacG glycosyltransferases participate in the elongation step forming the polyrhamnose backbone and capping the structure with unknown sugar residue. Polyrhamnose is transferred to the outer leaflet of the membrane presumably by the GacD/GacE ABC transporter. In the inner leaflet of the membrane, GacI aided by GacJ produces GlcNAc-P-Und [16] which then diffuses across the plasma membrane to the outer leaflet aided by GacK. Subsequently, GacL transfers GlcNAc to polyrhamnose using GlcNAc-P-Und as glycosyl donor [16]. GacH attaches GroP to the GlcNAc side-chains using phosphatidylglycerol as GroP donor [22]. Lastly, protein members of the LytR-CpsA-Psr phosphotransferase (LCP) family presumably attach GAC to peptidoglycan. Several details of this biosynthetic scheme are still speculative. But the overall organization is consistent with other isoprenol-mediated polysaccharide pathways.

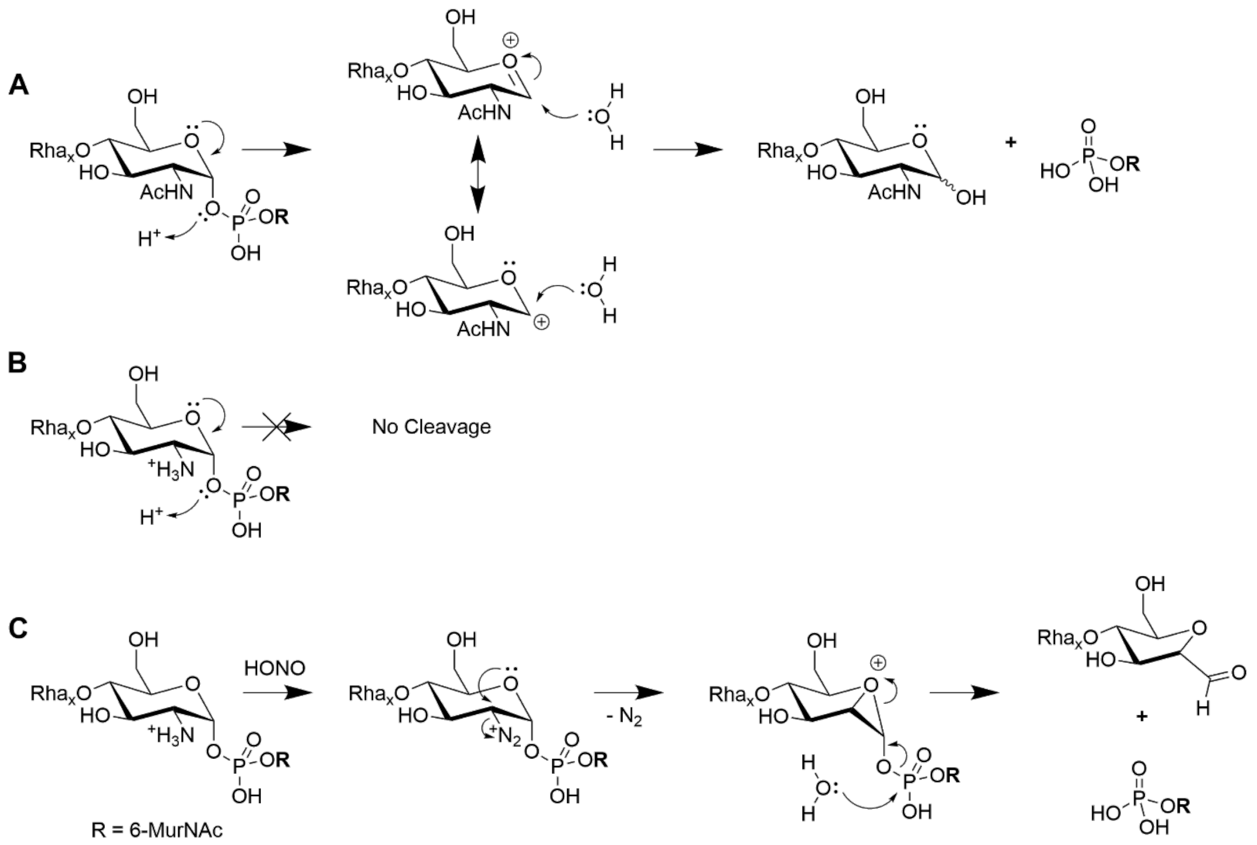

**Supplementary Fig. 2. Illustration of reaction mechanisms for hydrolysis of sugar 1-phosphates by 0.02 N HCl and deaminative cleavage of GlcN-1-phosphate by HONO.**
(**a**) Acid hydrolysis of GlcNAc 1-phosphate (and hexose 1-phosphates) proceeds efficiently by protonation of the O1-oxygen residue, facilitated by the resonance-stabilized oxonium ion intermediate. (**b**) The positively-charged GlcN-ammonium ion destabilizes the adjacent formation of the oxonium ion intermediate, preventing resonance stabilization of the intermediate. (**c**) Nitrous acid (HONO) forms a diazonium salt at the 2-position of GlcN; intramolecular attack of the GlcN ring oxygen on the diazonium center leads to a bicyclic oxonium intermediate and loss of nitrogen gas. Attack by water on the phosphodiester breaks the C1-O1 bond with concomitant opening of the strained three-membered oxonium ring, resulting in the formation of 2,5-anhydromannose and release of the 6-phosphate MurNAc monoester [26,27].

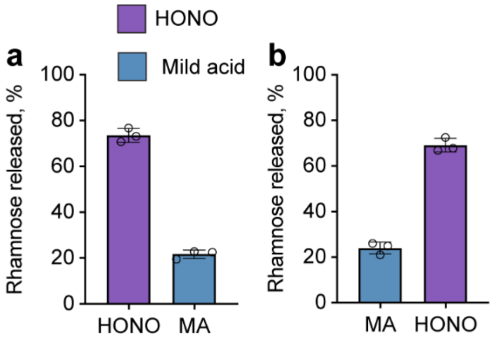

**Supplementary Fig. 3. Release of GAC from the GAS cell wall by sequential treatment with either mild acid or HONO.**

(**a**) GAC cell wall was subjected to HONO deamination, reisolated, and treated with mild acid. (**b**) GAC cell wall was subjected to mild acid hydrolysis, reisolated and deaminated with nitrous acid. Mild acid hydrolysis and nitrous acid deamination were conducted as described in Methods. The amount of GAC released from cell wall was estimated by the modified anthrone assay and normalized to total GAC content in cell wall. Symbols and error bars represent the mean and S.D. respectively (n = 3 biologically independent replicates).

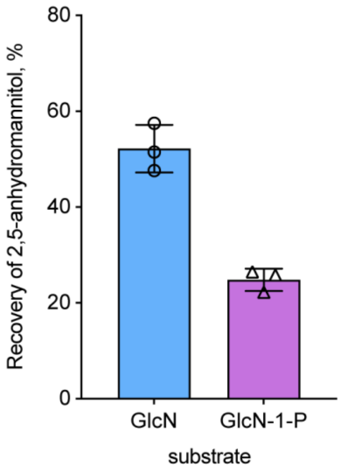

**Supplementary Fig. 4. Chemical yield of 2,5-anhydromannitol by HONO deamination of hexosamines.**

HONO deamination reactions contained 0.2 N Na acetate, pH 4.5, 1.5 M $NaNO_2$, and 400 nmol of either GlcN (left column) or GlcN 1-phosphate (right column) in a total volume of 0.01 mL. After 90 min at room temperature, the reactions were stopped by the addition of $NH_4OH$ and $NaBH_4$ to a final concentration of 1.5 M and 100 mg/mL, respectively. The reduction reaction was stopped after 2 h by the addition of acetic acid (5 μL) and the reactions were dried repeatedly out of methanol/0.1 % acetic acid under a stream of air to remove borate. The reactions were dissolved in water (344 μL) and an aliquot (5 μL) was taken for analysis. Mannitol (5 nmol, internal standard) was added and the aliquots were dried under air, peracetylated and analyzed by GC/MS as described in Methods for alditol acetate analysis. GlcN-1-phosphate concentration was verified by total phosphate analysis using the malachite green procedure. The amount of 2,5-anhydromannitol formed was calculated as described in Methods using authentic 2,5-anhydromannitol as standard. Symbols and error bars represent the mean and S.D. respectively (n = 3 biologically independent replicates).

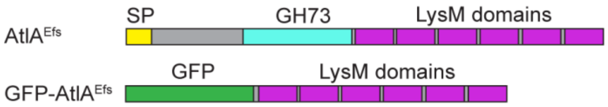

**Supplementary Fig. 5. The domain organization of AtlA$^{Efs}$ and the corresponding fluorescent fusion probe GFP-AtlA$^{Efs}$.** GH73 (cyan) denotes the catalytic domain; SP (yellow) indicates signal peptide.

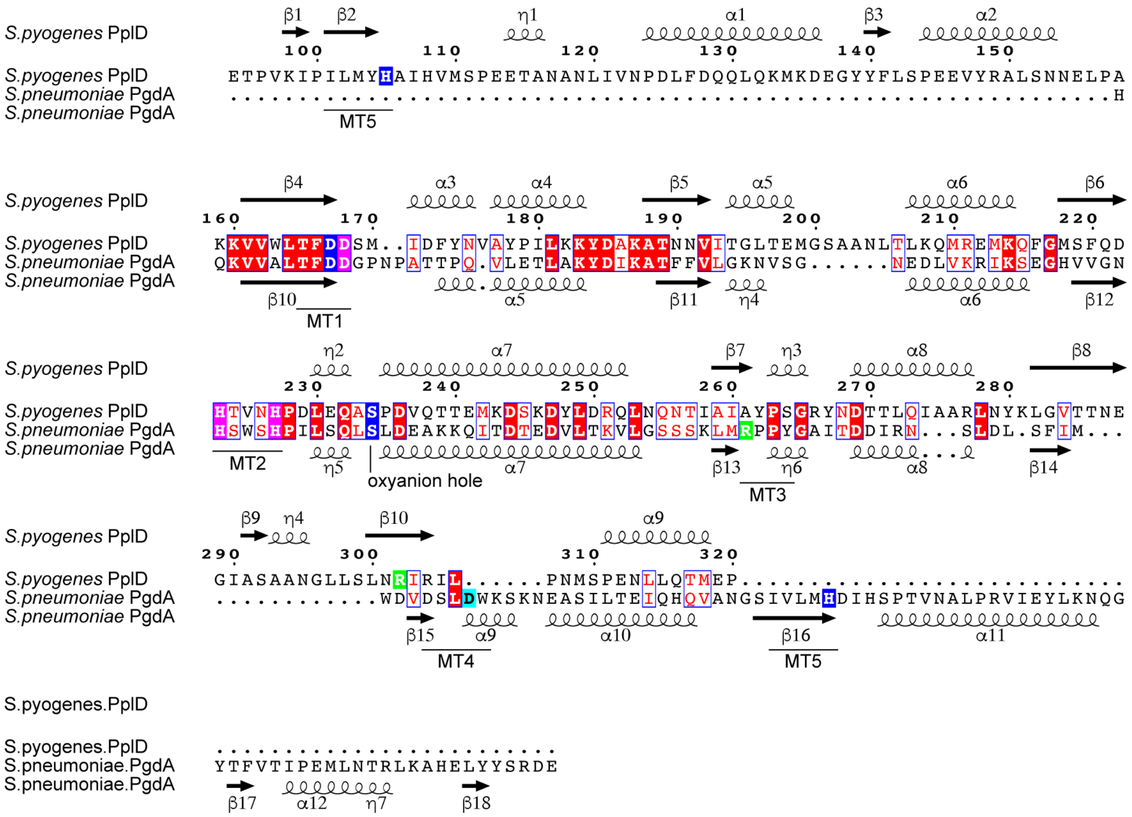

**Supplementary Fig. 6. Structure-based sequence alignment of extracellular domains of GAS PplD and *S. pneumoniae* PgdA.**

Residues implicated in the catalytic mechanism are highlighted in blue. The secondary structure elements are indicated above (PDB ID 6DQ3) and below (PDB ID 2C1G) [8] alignment.

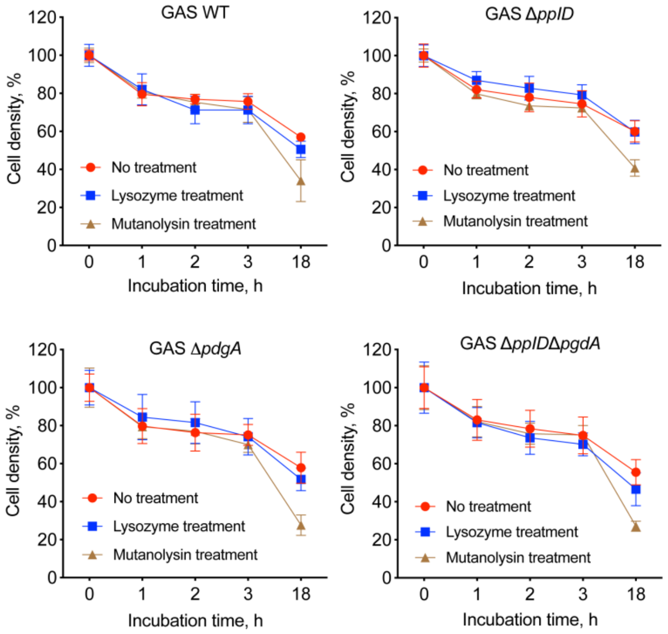

**Supplementary Fig. 7. Incubation of GAS WT, GASΔ*pplD*, GASΔ*pgdA*, GASΔ*pplD*Δ*pgdA*, with lysozyme.**

Bacterial strains were grown to an $OD_{600}$ of 0.5, centrifuged and resuspended in the same volume of sterile PBS with either 1 mg $mL^{-1}$ lysozyme, 62.5 U/ml mutanolysin (positive control) or no enzyme added (negative control). The bacterial suspension was incubated for 0, 1, 2, 3 and 18 h at 37°C. Lysis was monitored as the decrease in $OD_{600}$. Results were normalized to the $OD_{600}$ at time zero ($OD_{600}$ of 0.5). Data are mean values ± S.D., n = 3 biologically independent experiments.

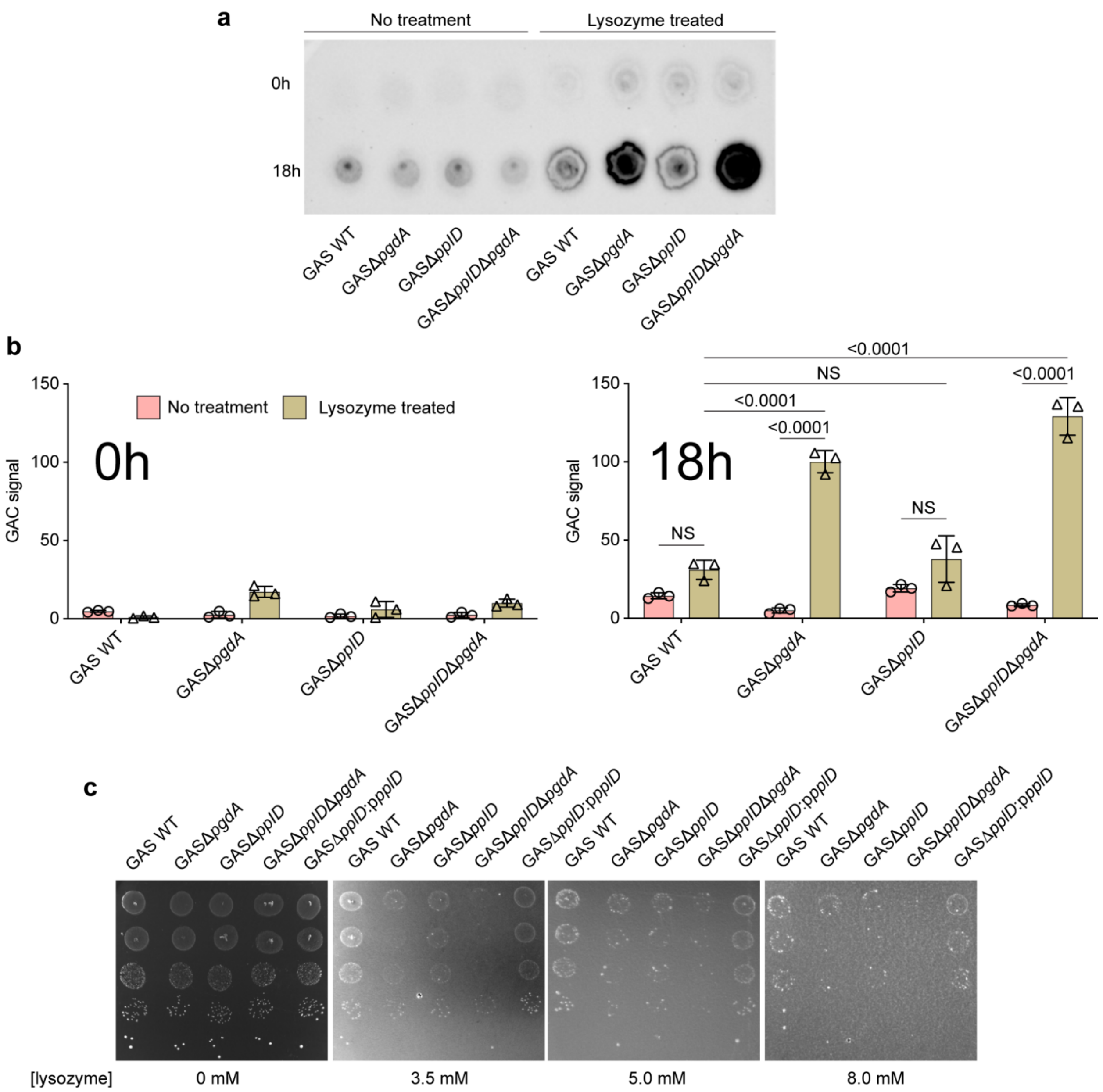

**Supplementary Fig. 8. Analysis of lysozyme sensitivity of GAS mutants deficient in *N*-deacetylases PgdA and PplD.**

(**a**) Dot-blot analysis for the water-soluble form of GAC released from the WT GAS, GASΔ*pplD*, GASΔ*pgdA* and GASΔ*pplD*Δ*pgdA* cells by lysozyme. GAS samples ($OD_{600}$ of 0.5) incubated for 0 and 18 h with or without lysozyme (1 mg $mL^{-1}$) were centrifuged (16,000 g, 3 min). The supernatant was concentrated using SpeedVac vacuum concentrator. 5 µl supernatant was spotted on a nitrocellulose membrane. GAC released to the supernatant was detected by anti-GAC antibodies as outlined in Methods. The experiment was performed independently three times and yielded the same results. Representative image from one experiment is shown. (**b**)

Immunoblot of the supernatant (representative image in Supplementary Fig. 8**a**) collected after 0 (left panel) and 18 h (right panel) of incubation was quantified using ImageJ. Data are mean values ± S.D., n = 3 biologically independent experiments. *P* values were calculated by two-way ANOVA with Tukey's multiple comparisons test. (**c**) Lysozyme sensitivity as tested in drop test assay using WT GAS, GASΔ*pplD*, GASΔ*pgdA*, GASΔ*pplD*Δ*pgdA* and GASΔ*pplD*:p*pplD*. Drop test assay experiment was performed at least three times.

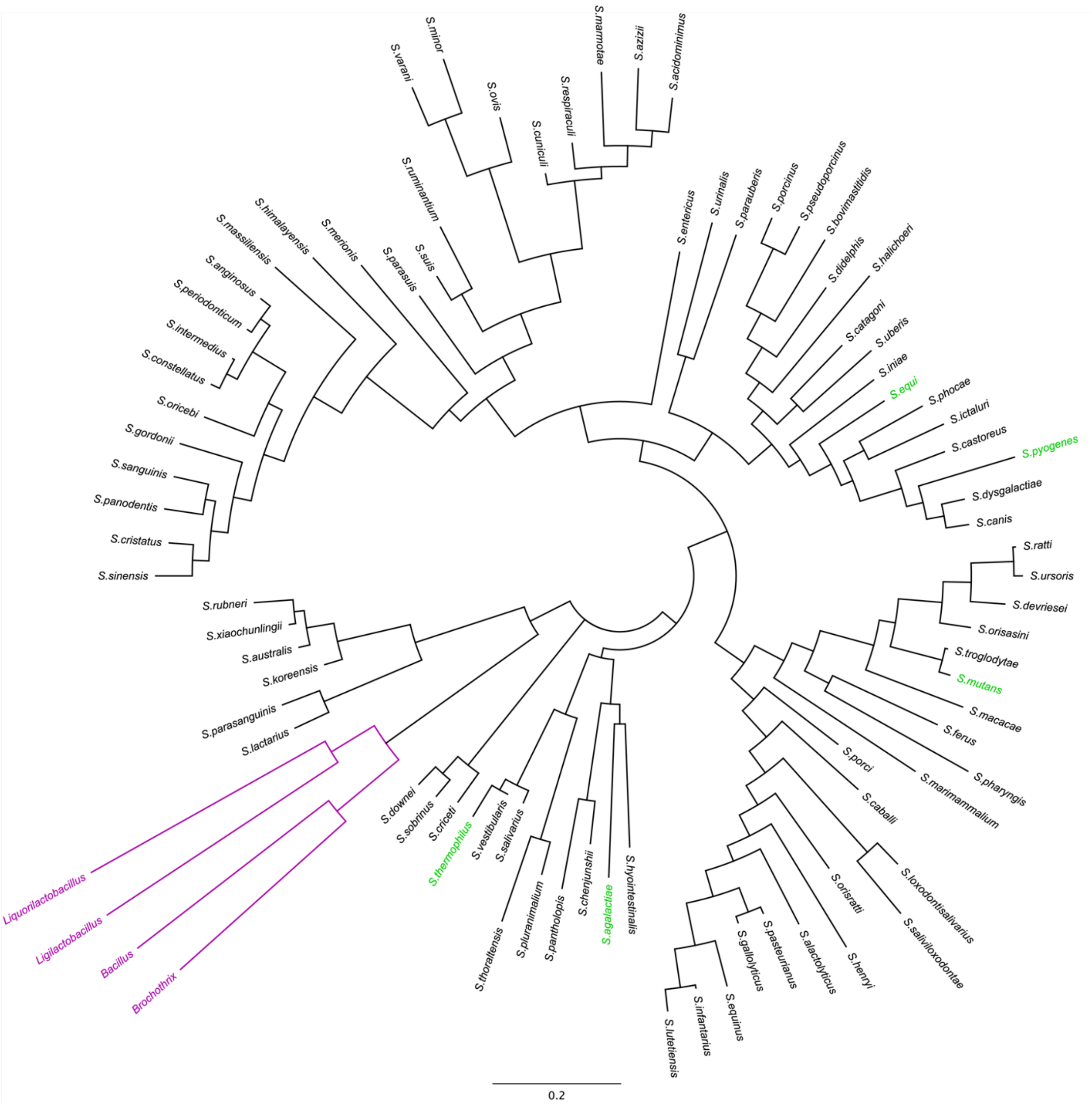

**Supplementary Fig. 9. Phylogenetic analysis of PplD homologs.**

A phylogenetic tree was generated using the predicted extracellular domains of 86 PplD homologs. The extracellular domains were predicted using the TMHMM server (http://www.cbs.dtu.dk/services/TMHMM/) [28]. There were a total of 234 amino acid positions in the final dataset. The Maximum Likelihood method and JTT matrix-based model [29] were used for evolutionary analysis of PplD homologs. The tree with the highest log likelihood (-16354.06) is shown. Initial tree(s) for the heuristic search were obtained automatically by applying Neighbor-Join and BioNJ algorithms to a matrix of pairwise distances estimated using the JTT model, and then selecting the topology with superior log likelihood value. The tree is drawn to scale, with branch lengths measured in the number of substitutions per site. Evolutionary analyses were conducted in MEGA X [30,31]. Streptococcal species are shown in black, non-streptococcal species are depicted in purple. *S. pyogenes*, *S. mutans*, *S. agalactiae*, *S. equi* and *S. thermophilus* are shown in green.

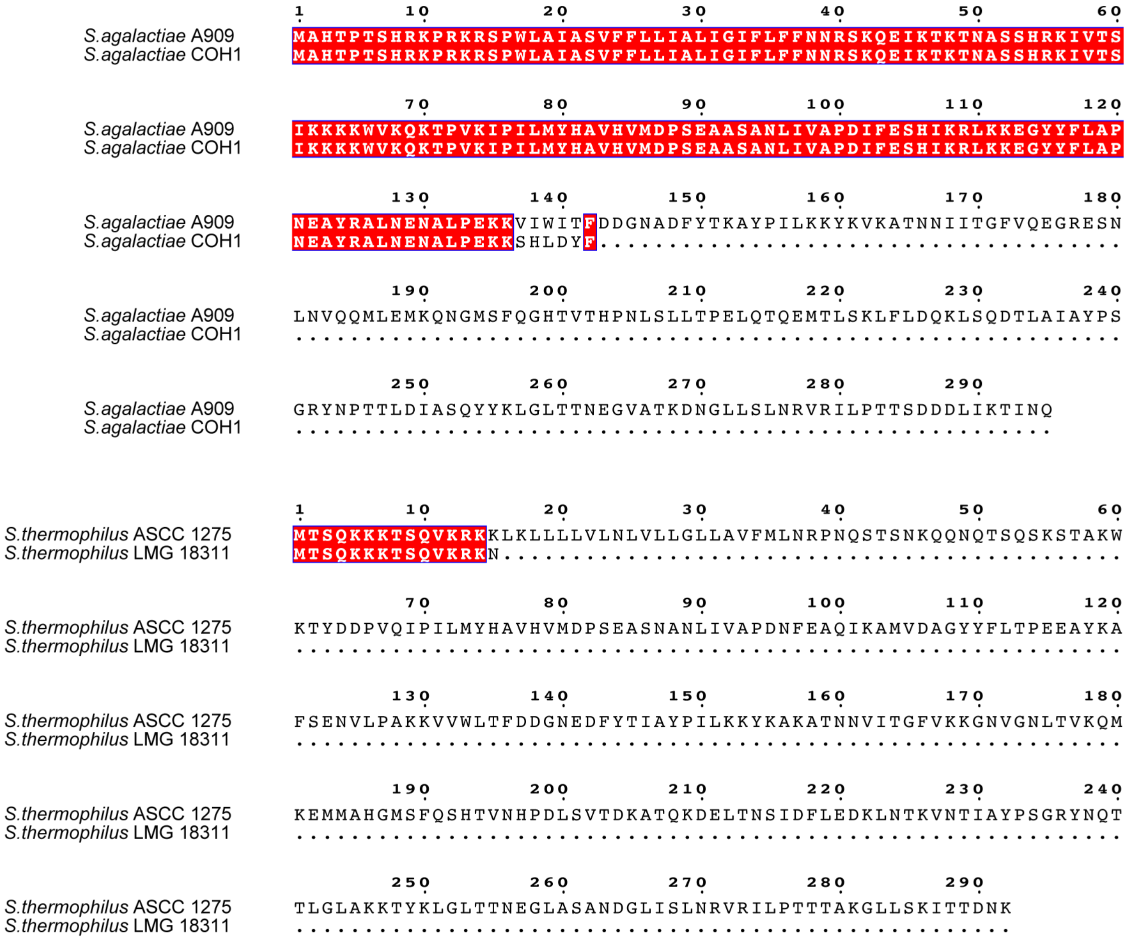

**Supplementary Fig. 10. Frame-shift mutations in *pplD* homologs result in truncated protein products in some streptococci strains.**
